# Cerebellar Purkinje cell stripe patterns reveal a differential vulnerability and resistance to cell loss during normal aging in mice

**DOI:** 10.1101/2025.01.26.634923

**Authors:** Sarah G. Donofrio, Cheryl Brandenburg, Amanda M. Brown, Tao Lin, Hsiang-Chih Lu, Roy V. Sillitoe

## Abstract

Age-related neurodegenerative diseases involve reduced cell numbers and impaired behavioral capacity. Neurodegeneration and behavioral deficits also occur during aging, and notably in the absence of disease. The cerebellum, which modulates movement and cognition, is susceptible to cell loss in both aging and disease. Here, we demonstrate that cerebellar Purkinje cell loss in aged mice is not spatially random but rather occurs in a pattern of parasagittal stripes. We also find that aged mice exhibit impaired motor coordination and more severe tremor compared to younger mice. However, the relationship between patterned Purkinje cell loss and motor dysfunction is not straightforward. Examination of postmortem samples of human cerebella from neurologically typical individuals supports the presence of selective loss of Purkinje cells during aging. These data reveal a spatiotemporal cellular substrate for aging in the cerebellum that may inform how neuronal vulnerability leads to neurodegeneration and the ensuing deterioration of behavior.

## INTRODUCTION

In addition to its well-known roles in motor function, the cerebellum is also involved in cognitive functions that include executive function, visuospatial memory, language, and emotional processing^1–11^. Accordingly, the cerebellum is a major culprit in movement disorders^12–27^, and it likely also contributes to autism spectrum disorders^28–34^, sleep disturbances^35–39^, and schizophrenia^40–42^. Despite the prevalence of cerebellar involvement in disease, the cerebellum is often overlooked in the context of aging. This omission has not gone unnoticed, with recent arguments for incorporating the cerebellum into our understanding of brain aging, which has historically focused on the cerebral cortex and hippocampus^43^. In support of this hypothesis, motor and cognitive behaviors are often impaired during normal aging. The decline in these behaviors could, in theory, be accompanied by cerebellar pathology, such as variations in cerebellar volume and alterations in its circuit connectivity^44–46^. Therefore, the cerebellum is a potentially critical model system for better understanding structure, function, and behavior of the brain during aging.

Although neurodegeneration is associated with disease pathogenesis, cerebellar degeneration can also occur during aging in the absence of disease. Elderly patients have decreased cerebellar volume compared to young patients^47–50^, and longitudinal studies have shown decreased cerebellar volume in healthy older individuals over time^51–55^. At the cellular level, otherwise healthy aged patients have significantly decreased Purkinje cell density^56–58^. Depending on severity, age-related cerebellar atrophy can have detrimental and often debilitating effects on motor function. For example, decreased cerebellar volume in the elderly is correlated with impaired eyeblink conditioning, a cerebellum-dependent associative learning task, as well as slower gait and impaired balance, two aspects of motor function that involve the cerebellum^59,60^. Furthermore, there are reported structure-function relationships between the volume of specific cerebellar regions and the performance of sensorimotor tasks in young and old participants^61^. These results suggest that cerebellar atrophy is associated with deficits in cerebellum-related functions during normal aging. However, the cellular nature of this structure-function relationship is still poorly understood.

Similar to humans, control mice and rats also experience Purkinje cell loss during aging^62–71^, as well as motor dysfunction. At around 12 months of age, mice begin to display impaired performance on the rotarod task, a test of motor coordination and motor learning^72–74^, as well as impaired eyeblink conditioning^73^, now a classical test for Pavlovian learning. Interestingly, there is evidence to suggest that mice have significantly decreased Purkinje cell numbers starting at 18 months, corresponding with impaired delay eyeblink conditioning^75^. These studies suggest that cerebellum-associated motor function declines with age in mice and that this decline may be accompanied by Purkinje cell loss. However, the precise regionality of the cerebellum was not considered in these studies, and we argue that age-related changes follow a fundamental scheme.

Compared to the patterned Purkinje cell loss reported in different disease models, relatively little is known about how Purkinje cell loss affects different regions of the cerebellum during normal aging. Neuroimaging in humans has revealed that cerebellar volume in different lobules can be differentially affected by aging^47,55,61,76–79^. However, a closer examination of any region-specific cellular differences is lacking. Other studies have found minimal regional differences beyond the observation that most age-related Purkinje cell loss seems to occur in the anterior cerebellum^56,57^. In fact, multiple studies have found that Purkinje cell loss is uniform across the latero-lateral extent of the cerebellar cortex^62–66,69^. However, cerebellar organization and patterning are much more complex than the broad medio-lateral and anterior-posterior differences can account for.

The cerebellum is highly compartmentalized on multiple levels as determined by genetics and developmental, anatomical, and electrophysiological studies. Based on development and gene expression boundaries, the cerebellum can be divided into transverse zones: the anterior (lobules I-V), central (lobules VI and VII), posterior (lobule VIII and anterior lobule IX), and nodular zones (posterior lobule IX and lobule X)^80^. Within each transverse zone, subpopulations of Purkinje cells are divided into stripes based on gene expression patterns, which differ between transverse zones^81–87^. Numerous stripe markers exist, with overlapping, partially overlapping, or unique expression patterns. The most well-studied stripe marker is zebrin II^88–90^, the expression pattern of which is remarkably consistent from animal to animal and conserved across mammals^91,92^. The identity of individual Purkinje cells, which can express different combinations of patterned markers, is established when each Purkinje cell is born^93–96^, although the exact relationship between developmental Purkinje cell clusters and adult stripes has been only partially resolved^97^. The cerebellar stripe patterns represent one component of a greater network architecture. For example, longitudinal groups of Purkinje cells are related in the rostrocaudal axis based on specific inputs from the inferior olive through climbing fibers and specific outputs to the cerebellar nuclei through the Purkinje cell axons^98^. Each longitudinal zone together with its efferent and afferent pathways comprises a functional unit referred to as a cerebellar module^99^. Based on these overlapping levels of organization, the cerebellum is predicted to be composed of hundreds or thousands of modules. This hypothesis posits that the multiple maps of cerebellar compartmentation that are visualized with different approaches comprise a single overarching map, with alignment between zones, stripes, and modules^98^. Here, we wondered whether the patterns that are revealed by stripe markers reflect a map of Purkinje cell degeneration and eventual cell loss with age and if there is a functional significance to this patterned cellular demise.

To test for and investigate the potential contribution of patterned Purkinje cell loss to aging in mice, we used a combination of wholemount immunohistochemistry^100,101^ and analysis of Purkinje cell patterns on histological tissue slices. These techniques enabled us to visualize Purkinje cells across the surfaces of entire cerebella of normal aged mice and examine their cellular level changes, respectively. We also used genetically driven fluorescent reporter labeling to specifically label Purkinje cells. Using these techniques, we uncovered that some, but not all, aged mice have Purkinje cell loss that occurs in a pattern of parasagittal stripes. In addition, upon immunostaining coronal sections of cerebellar tissue, we found that the pattern of age-related Purkinje cell loss is unique compared to the most common patterns of Purkinje cell loss that have been reported in numerous mouse models of disease. Behavioral tests revealed deficits in motor function in aged mice compared to young mice. Finally, we observed patches of Purkinje cell degeneration in postmortem tissue obtained from neurologically normal humans. We therefore have three key findings to report in this series of studies: 1) Purkinje cell degeneration and cell loss is a prominent feature of the otherwise healthy aging mouse cerebellum, 2) Purkinje cell loss during normal aging in mice does not occur in a random manner and instead occurs in an array of parasagittal stripes that reflect the normal developmental, anatomical, and functional topography of the mammalian cerebellum, and 3) this mode of patterned cell loss may be conserved and reflect a fundamental heterogeneity of the human cerebellum that distinguishes cells that are vulnerable versus resistant to deterioration in health and disease. Together, these data underscore the potential clinical significance of our findings of patterned Purkinje cell degeneration and loss in normal aging mice, as they may inform on the design and development of effective therapies for specific neurological and neuropsychiatric conditions that are defined by compromised cerebellar structure and function.

## RESULTS

Adult-onset neurodegeneration typically impacts specific regions of the brain. For instance, in diseases such as Alzheimer’s disease and Parkinson’s disease, neurodegeneration affects the cortex and basal ganglia, resulting in defective cognitive versus motor circuits, respectively. A similar phenomenon occurs during normal aging, as certain brain regions are more susceptible to degeneration^102^. Interestingly, the cerebellum is a target in age-related degeneration, and its cellular demise can lead to both cognitive and motor impairments. We examine whether different populations of cells are more susceptible than others. What dictates this regional vulnerability and whether these processes and sensitive neuronal subpopulations are the same across different diseases and typical aging is unknown. To begin to address this problem, here we sought to test whether cerebellar Purkinje cell loss follows a region-specific pattern during normal aging.

### Aged mice have Purkinje cell loss that occurs in parasagittal stripes

We started by asking whether normal mice exhibit patterned Purkinje cell loss during aging. Previous studies have relied on tissue sections alone for identifying the presence of degeneration and cell loss, although studying the complexity of the cerebellum on individual slices limits one’s ability to visualize topographic patterns. In our study, to test whether age-related Purkinje cell loss is indeed patterned, we examined a total of 52 cerebella from aged mice (between 13 and 25 months of age; Table 1). We used a combination of techniques to visualize Purkinje cells, including wholemount immunohistochemistry^100,101^ with calbindin antibodies (n=5), wholemount and light sheet imaging of a Purkinje cell-specific reporter in transgenic mice (n=21 and n=2, respectively), and histology on coronal tissue sections (n=34; Table 1). Using these techniques, we observed that Purkinje cell loss across the cerebella of aged mice is not uniform, as has been previously reported^62–66,69^, but forms a pattern of Purkinje cell loss. Importantly, the pattern was composed of parasagittal stripes that are symmetrical about the midline. In the calbindin-labeled wholemount cerebella, the stripes appeared as alternating dark stripes of surviving Purkinje cells and light stripes where Purkinje cells have presumably degenerated (Fig. 1A). The stripes were visible in the anterior, central, and posterior zones of the cerebellum, as well as in the paraflocculi. Closer inspection of the wholemount cerebella revealed the shapes of individual Purkinje cells within the dark stripes, with the dendrites and cell bodies clearly visible (Fig. 1B). Purkinje cell axons with torpedoes, a pathological sign of Purkinje cell neurodegeneration in disease^103^ and normal aging^104^, were also observed in wholemount cerebella (Fig. 1B). The presence of the axonal pathology suggests that the observed Purkinje cell loss in aged mice may be due to and potentially accompanied by a process of neurodegeneration that causes cell loss over time.

**Figure 1:**
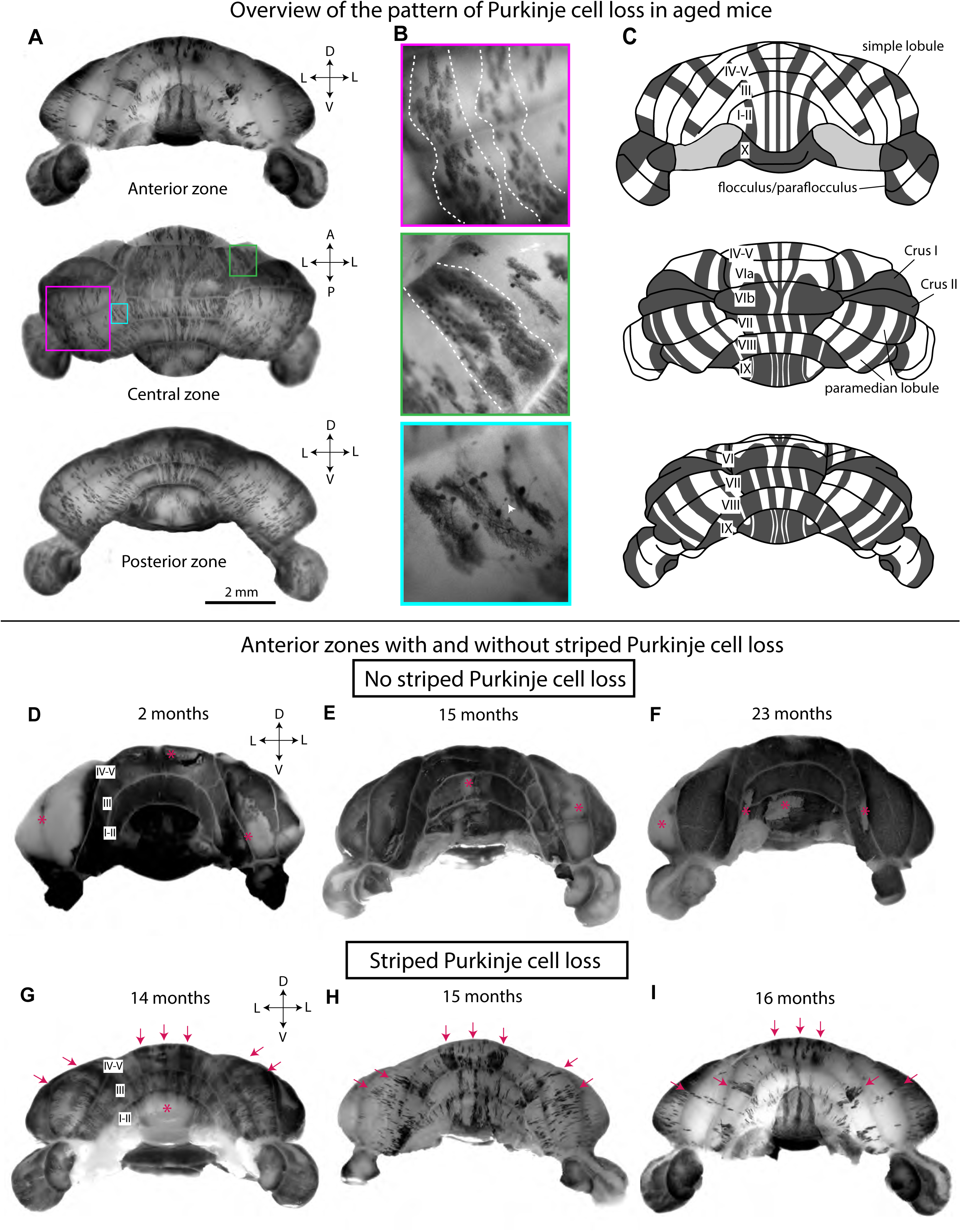
Wholemount immunohistochemistry of the cerebellum reveals striped Purkinje cell loss across the cerebellar cortex of some aged mice. A) Wholemount cerebellum of a 16-month-old mouse immunostained for calbindin and viewed from different angles. D = dorsal; L = lateral; V = ventral; A = anterior; P = posterior. Scale bar = 2 mm. B) High-magnification images of Purkinje cells in the wholemount cerebellum of an aged mouse. Dotted lines indicate stripes of surviving Purkinje cells, and the white arrowhead indicates an axonal torpedo. C) Schematic of the pattern of age-related Purkinje cell loss based on wholemount cerebella, where dark gray stripes represent bands largely composed of surviving Purkinje cells and white stripes represent bands where most Purkinje cells have degenerated. Cerebellar lobules are labeled with Roman numerals. D-I) Wholemount cerebella of mice immuno-stained for calbindin and viewed from the anterior zone: D) 2-month-old mouse; E) 15-month-old mouse without Purkinje cell loss; F) 23-month-old mouse without Purkinje cell loss; G) 14-month-old mouse with striped Purkinje cell loss; H) 15-month-old mouse with striped Purkinje cell loss; I) 16-month-old mouse with striped Purkinje cell loss. Asterisks indicate staining artifacts, either caused by continuous rubbing of the cerebellum during staining (hemispheres in panels D-F and lobules I-II in panel G) or accidental removal of surface tissue during dissection (lobules I-II in panel F). Arrows indicate bands of surviving Purkinje cells that are consistent across mice with striped Purkinje cell loss. D = dorsal; L = lateral; V = ventral. Cerebellar lobules are labeled with Roman numerals.

**Table 1:** Information from mice used in this study. Mice from the same litter are indicated with the same letter.

| Sex | Age (months) | PC loss | Genotype | Littermates | Technique |
| --- | --- | --- | --- | --- | --- |
| M | 7 | striped PC loss | *Pcp2*<sup>Cre+/-</sup>; *Ai40D* | a | GFP wholemount |
| F | 7.2 | no PC loss | *Pcp2*<sup>Cre+/-</sup>; *Ai40D* |  | GFP wholemount |
| M | 7.4 | no PC loss | no mutant alleles | a | coronal sections |
| F | 11.7 | striped PC loss | *Pcp2*<sup>Cre+/-</sup>; *Ai40D* | b | GFP wholemount |
| M | 13.7 | no PC loss | *Ai32*<sup>+/+</sup> |  | coronal sections |
| M | 13.9 | striped PC loss | no mutant alleles | c | coronal sections |
| M | 14.3 | non-striped PC loss | *Pcp2*<sup>Cre+/-</sup>; *Ai32* |  | GFP wholemount |
| M | 14.3 | striped PC loss | *Pcp2*<sup>Cre+/-</sup>; *Ai32*<sup>+/+</sup> | d | GFP wholemount |
| M | 14.4 | no PC loss | *Pcp2*<sup>Cre+/-</sup>; *Ai32*<sup>+/+</sup> | d | GFP wholemount |
| F | 14.5 | striped PC loss | *Pcp2*<sup>Cre+/-</sup>; *Ai32*<sup>+/-</sup> |  | GFP wholemount |
| M | 14.7 | striped PC loss | *Tau*<sup>Isl-mGFP-lacZ</sup> |  | calbindin wholemount |
| F | 15.1 | striped PC loss | *Pcp2*<sup>Cre+/-</sup>; *Ai32* |  | GFP wholemount |
| M | 15.3 | striped PC loss | *Pcp2*<sup>Cre+/-</sup>; *Tau*<sup>Isl-mGFP-lacZ</sup> | e | calbindin wholemount |
| F | 15.3 | no PC loss | *Tau*<sup>Isl-mGFP-lacZ</sup>; *ROSA26*<sup>Isl-DTR+/-</sup> | e | calbindin wholemount |
| M | 15.7 | non-striped PC loss | *Pdx1*<sup>Cre+/-</sup> |  | coronal sections |
| F | 16.3 | striped PC loss | *Tau*<sup>Isl-mGFP-lacZ</sup>; *ROSA26*<sup>Isl-DTR+/-</sup> |  | calbindin wholemount |
| F | 16.4 | no PC loss | *Tau*<sup>Isl-mGFP-lacZ</sup> |  | coronal sections |
| F | 16.4 | striped PC loss | *Pcp2*<sup>Cre+/-</sup>; *Ai40D*<sup>+/-</sup> | f | GFP wholemount, coronal sections |
| M | 16.5 | non-striped PC loss | *Ai32*<sup>+/-</sup> | g | coronal sections |
| M | 16.5 | no PC loss | *Gdnf*<sup>CreER+/-</sup>; *Ai32*<sup>+/-</sup> | g | coronal sections |
| M | 16.5 | non-striped PC loss | *Ai32*<sup>+/-</sup> | g | coronal sections |
| M | 16.6 | no PC loss | *Tau*<sup>Isl-mGFP-lacZ</sup>; *ROSA26*<sup>Isl-DTR+/-</sup> |  | coronal sections |
| F | 16.8 | striped PC loss | *Pcp2*<sup>Cre+/-</sup>; *Ai40D* |  | GFP wholemount, coronal sections |
| M | 16.8 | no PC loss | no mutant alleles | h | coronal sections |
| M | 16.8 | no PC loss | no mutant alleles | h | coronal sections |
| M | 16.9 | no PC loss | *Pcp2*<sup>Cre+/-</sup> | i | coronal sections |
| M | 16.9 | no PC loss | *Tau*<sup>Isl-mGFP-lacZ</sup>; *ROSA26*<sup>Isl-DTR+/-</sup> | i | coronal sections |
| M | 16.9 | no PC loss | no mutant alleles | j | coronal sections |
| M | 16.9 | no PC loss | no mutant alleles | j | coronal sections |
| M | 17 | striped PC loss | *Pcp2*<sup>Cre+/-</sup>; *Ai40D*<sup>+/-</sup> | k | GFP wholemount, light sheet |
| M | 17.2 | striped PC loss | *ROSA26*<sup>Isl-DTR+/-</sup> | c | coronal sections |
| M | 17.3 | no PC loss | *Ai40D*<sup>+/-</sup> | k | GFP wholemount, coronal sections |
| M | 17.4 | striped PC loss | *Pcp2*<sup>Cre+/-</sup>; *Ai40D*<sup>+/-</sup> | k | GFP wholemount |
| M | 17.4 | no PC loss | *Ai32*<sup>+/+</sup>; *VGAT*<sup>+/fx</sup> |  | coronal sections |
| F | 18 | striped PC loss | *Pcp2*<sup>Cre+/-</sup>; *Ai32* | b | GFP wholemount |
| M | 18.3 | no PC loss | *Mash1*<sup>CreER+/-</sup>; *ROSA26*<sup>Isl-eYFP</sup>; *VGAT*<sup>fx/fx</sup> |  | coronal sections |
| F | 18.3 | striped PC loss | *Pcp2*<sup>Cre+/-</sup>; *Ai40D*<sup>+/-</sup> | l | GFP wholemount |
| F | 18.5 | striped PC loss | *Pcp2*<sup>Cre+/-</sup>; *Ai32* | m | coronal sections |
| F | 18.9 | no PC loss | *Ai40D*<sup>+/-</sup> | l | coronal sections |
| F | 19.8 | non-striped PC loss | *ROSA26*<sup>Isl-eYFP</sup>; *VGAT*<sup>fx/fx</sup> |  | coronal sections |
| F | 20.1 | non-striped PC loss | *ROSA26*<sup>Isl-eYFP</sup>; *VGAT*<sup>fx/fx</sup> | n | coronal sections |
| F | 20.1 | non-striped PC loss | *ROSA26*<sup>Isl-eYFP</sup>; *VGAT*<sup>fx/fx</sup> | n | coronal sections |
| F | 20.9 | striped PC loss | *Pcp2*<sup>Cre+/-</sup>; *Ai32* | m | coronal sections |
| F | 20.9 | striped PC loss | *Pcp2*<sup>Cre+/-</sup>; *Ai32* | m | coronal sections |
| M | 20.9 | striped PC loss | *Pcp2*<sup>Cre+/-</sup>; *Ai32* | m | coronal sections |
| M | 21.2 | striped PC loss | *Pcp2*<sup>Cre+/-</sup>; *Ai32* | b | GFP wholemount, coronal sections |
| M | 21.2 | striped PC loss | *Pcp2*<sup>Cre+/-</sup>; *Ai32* | b | GFP wholemount, coronal sections |
| M | 21.4 | striped PC loss | *Pcp2*<sup>Cre+/-</sup>; *Ai40D* | o | GFP wholemount, coronal sections |
| M | 21.4 | striped PC loss | *Pcp2*<sup>Cre+/-</sup>; *Ai40D* | o | GFP wholemount, light sheet |
| F | 23.3 | no PC loss | *VGAT*<sup>fx/fx</sup> |  | calbindin wholemount |
| F | 25.5 | striped PC loss | *Pcp2*<sup>Cre+/-</sup>; *Ai40D*<sup>+/-</sup> | f | GFP wholemount, coronal sections |
| M | 25.5 | striped PC loss | *Pcp2*<sup>Cre+/-</sup>; *Ai40D* | f | GFP wholemount, coronal sections |

We observed considerable variability in terms of Purkinje cell loss in the aged mice. Some aged mice displayed a lack of Purkinje cell loss comparable to young control mice, even at 23 months of age (Fig. 1D-F). Of the 52 aged cerebella examined, 19 lacked appreciable Purkinje cell loss, 26 had clearly striped Purkinje cell loss, and 7 had Purkinje cell loss that did not appear striped (Table 1; Supplementary Fig. 1). This inconsistency in the loss of Purkinje cells is not due solely to relative age, as a 14-month-old cerebellum had striped Purkinje cell loss (Fig. 1G), whereas a 23-month-old cerebellum did not (Fig. 1F; Table 1). Despite this, when striped Purkinje cell loss did occur in the aged mice, the neurodegeneration appears progressive, meaning that older mice were more likely to display more widespread Purkinje cell loss compared to younger mice (Fig. 1G-I). Aged mice with striped Purkinje cell loss consistently displayed the same pattern of neurodegeneration (Fig. 1G-I). Therefore, Purkinje cells that are more susceptible to age-related neurodegeneration may belong to the same subpopulation of neurons–and have the same identity– across different mice.

Of the 52 aged mice whose cerebella were examined, 32 were male and 20 were female. We found that 12 of the 20 aged female mice and 14 of the 32 aged male mice had striped Purkinje cell loss (Table 1, Supplementary Fig. 1A). Analysis of the data with Fisher’s exact test revealed that the presence of striped Purkinje cell loss is not sex dependent. However, a greater percentage of the aged females displayed Purkinje cell loss compared to the aged males (Supplementary Fig. 1A), suggesting that females may be more susceptible to Purkinje cell loss during aging. Overall, these results suggest that there is considerable variability plus context specificity in the spatiotemporal features of Purkinje cell degeneration, eventual Purkinje cell loss and their associated pattern among aged mice.

Interestingly, among the mice born in the same litter, we found that one littermate could have striped Purkinje cell loss while the other littermate did not (Fig. 1E and H). This observation was especially evident as observed on surface mapping using the wholemount immunohistochemical staining approach (Fig. 1). This difference in cell loss was observed in five sets of littermates, with each set from a different litter (Table 1). This data suggests that even genetically similar mice raised in the same cage can display dramatic differences in age-related neurodegeneration.

Taken together, these findings suggest that 1) Purkinje cell subpopulations are differentially vulnerable to death during normal aging; 2) the loss of Purkinje cells, according to age-related vulnerability, can result specifically in a striking pattern of parasagittal stripes; and 3) the presence or absence of striped Purkinje cell loss in aged mice is not driven solely by sex or relative age.

### Age-related Purkinje cell loss is due to neurodegeneration and the loss of cells

In mouse models with neurodegenerative ataxia, calbindin and other Purkinje cell-specific genes and proteins are downregulated prior to the onset of motor dysfunction and Purkinje cell loss^105,106^. This molecular signature raises the possibility that the indication of Purkinje cell loss revealed by antibody staining may in fact be the result of reduced Purkinje cell marker expression rather than neurodegeneration. To distinguish between these possibilities, we tested whether Purkinje cells in aged mice were degenerating. First, we observed common hallmarks of Purkinje cell neurodegeneration, such as thickened axons, axonal torpedoes, and shrunken dendritic arbors, in aged mice with striped Purkinje cell loss (n=8) but not in young mice (n=5; Fig. 2A and B). Quantification of molecular layer thickness in lobule VIII revealed that aged mice with striped Purkinje cell loss have significantly thinner molecular layers compared to young mice (Fig. 2C), likely due to the regressed dendrites of degenerating Purkinje cells. Aged mice without Purkinje cell loss have minimal axonal pathology (Fig. 2A), and the thickness of the molecular layer is comparable to that of young mice (Fig. 2B and C), suggesting that in aged mice without Purkinje cell loss, Purkinje cells are not undergoing major neurodegeneration. We also confirmed whether the pattern of Purkinje cell loss was the same using two different calbindin antibodies (young n=4, aged n=6; Supplementary Fig. 2).

**Figure 2:**
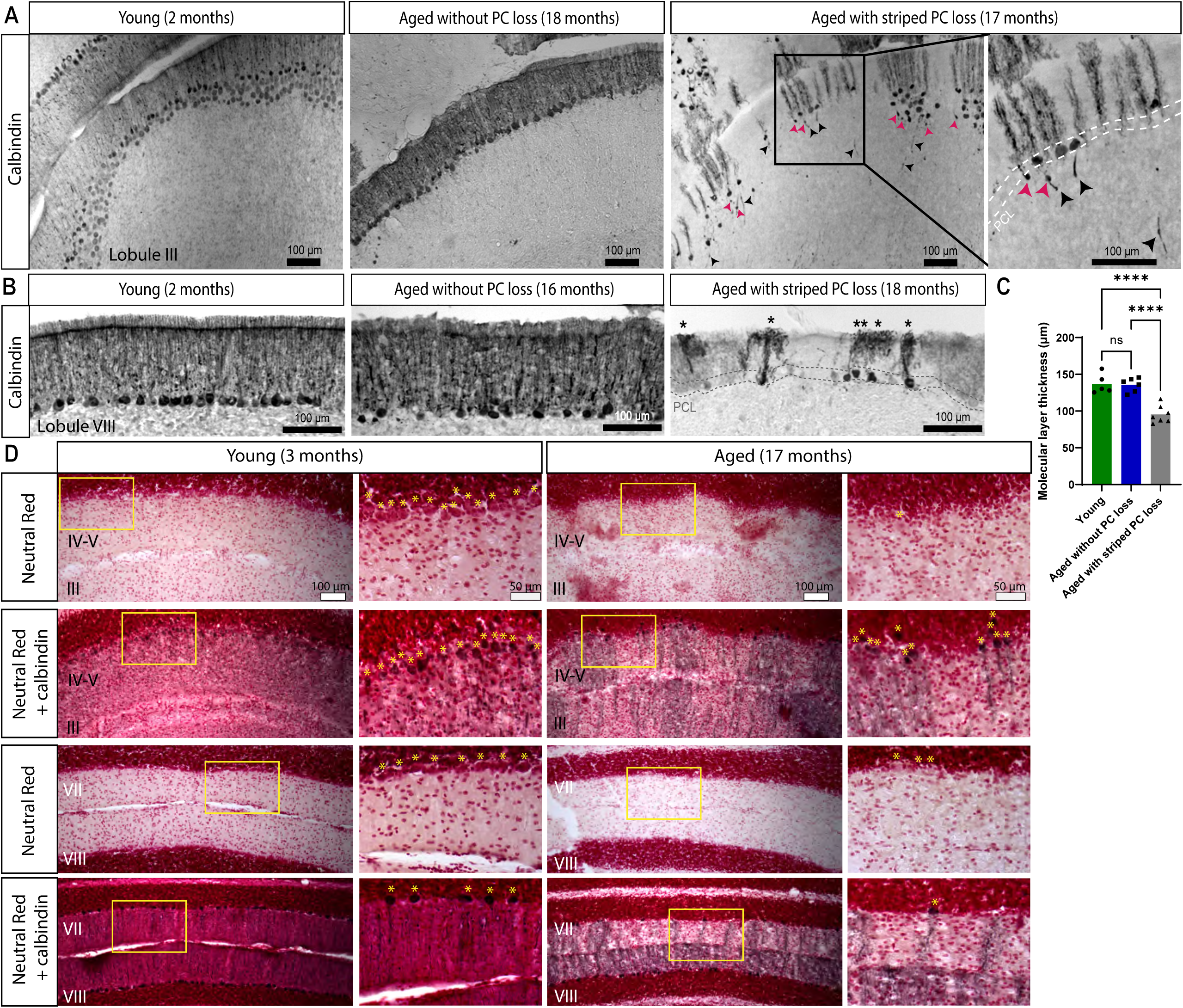
Aged mice with striped Purkinje cell loss show cellular level anatomical hallmarks of degeneration. A) Coronal cut cerebellar tissue sections of lobule III immunostained for calbindin. Black arrowheads indicate thickened axons, and pink arrowheads indicate axonal torpedoes. Dashed lines indicate the Purkinje cell layer (PCL). Scale bar = 100 μm. B) Coronal cut cerebellar tissue sections of lobule VIII immunostained for calbindin. Asterisks indicate shrunken dendritic arbors. Scale bar = 100 μm. C) Quantification of molecular layer thickness in lobule VIII; **** indicates p ≤ 0.0001. D) Coronal cut cerebellar tissue sections either stained with Neutral Red or immunostained for calbindin and stained with Neutral Red. Asterisks indicate Purkinje cell bodies. Scale bar = 100 μm; inset scale bar = 50 μm.

Second, to verify that the loss of Purkinje cells was not due to downregulation of calbindin specifically, we used Neutral Red, a dye that stains lysosomes, to label surviving cells in cerebellar tissue^107^. Adjacent tissue sections were immunostained for calbindin and/or stained with Neutral Red to locate regions with striped Purkinje cell loss. In the aged mice (n=6), regions of the Purkinje cell and molecular layers without calbindin staining, which indicated missing Purkinje cells, also lacked Neutral Red-positive cell bodies, which based on their morphology were easily distinguished from other cell types by their size and position in young mice (n=6; Fig. 2D).

Third, to verify the presence of striped Purkinje cell degeneration during normal aging, we used adult Purkinje cell-specific fluorescent reporter mice, which express a fluorescent reporter specifically in all Purkinje cells because the reporter is driven by *Pcp2^Cre^*. These mice present two advantages for visualizing age-related Purkinje cell loss: 1) Cre expression begins on E17 and continues until all Purkinje cells express Cre in adulthood^108^, meaning that even if *Pcp2* gene and protein expression are downregulated in advanced age^109^ or prior to neurodegeneration^106^, reporter expression will remain constant as it would have already been activated in all Purkinje cells and its perdurance would allow continued marking of the recombined cells; and 2) Purkinje cell-specific reporter expression allows for the visualization of Purkinje cells without relying on calbindin, which can be downregulated with advanced age^109–111^ or prior to neurodegeneration^105,106^. We found that the cerebella of young Purkinje cell-specific fluorescent reporter mice (n=4) displayed uniform reporter expression across the surface, whereas the cerebella of aged Purkinje cell-specific fluorescent reporter mice (n=17) displayed reporter expression in alternating parasagittal stripes of greater and lesser intensity (Fig. 3A and B). Furthermore, middle-aged mice (11-15 months old; n=4) tended to have less pronounced bands compared to mice 16 months and older, which had clearer, more widespread bands (Fig. 3A and B), suggesting a progression of Purkinje cell loss with advanced age. The cerebella of middle-aged mice have striped reporter expression in the anterior zone, large regions with reduced reporter expression in the medial vermis and paravermis, and alternating bands in lobule VIII (Fig. 3A and B), suggesting that age-related Purkinje cell loss may begin in these regions before spreading with advancing age. To confirm that the Purkinje cell-specific reporter expression and the calbindin wholemount staining reflected the same pattern, we co-stained coronal cerebellar tissue sections of Purkinje cell-specific fluorescent reporter mice for calbindin and GFP. In young mice, both calbindin and GFP were expressed in all Purkinje cells (n=3), and in aged mice, calbindin and GFP were expressed in identical, overlapping stripes (n=5; Fig. 3D and Supplementary Fig. 3) that reflected the pattern of age-related Purkinje cell loss. These data indicate that the pattern of age-related Purkinje cell loss we revealed with calbindin wholemount staining matches the pattern that we next observed with reporter expression in Purkinje cell-specific fluorescent reporter mice.

**Figure 3:**
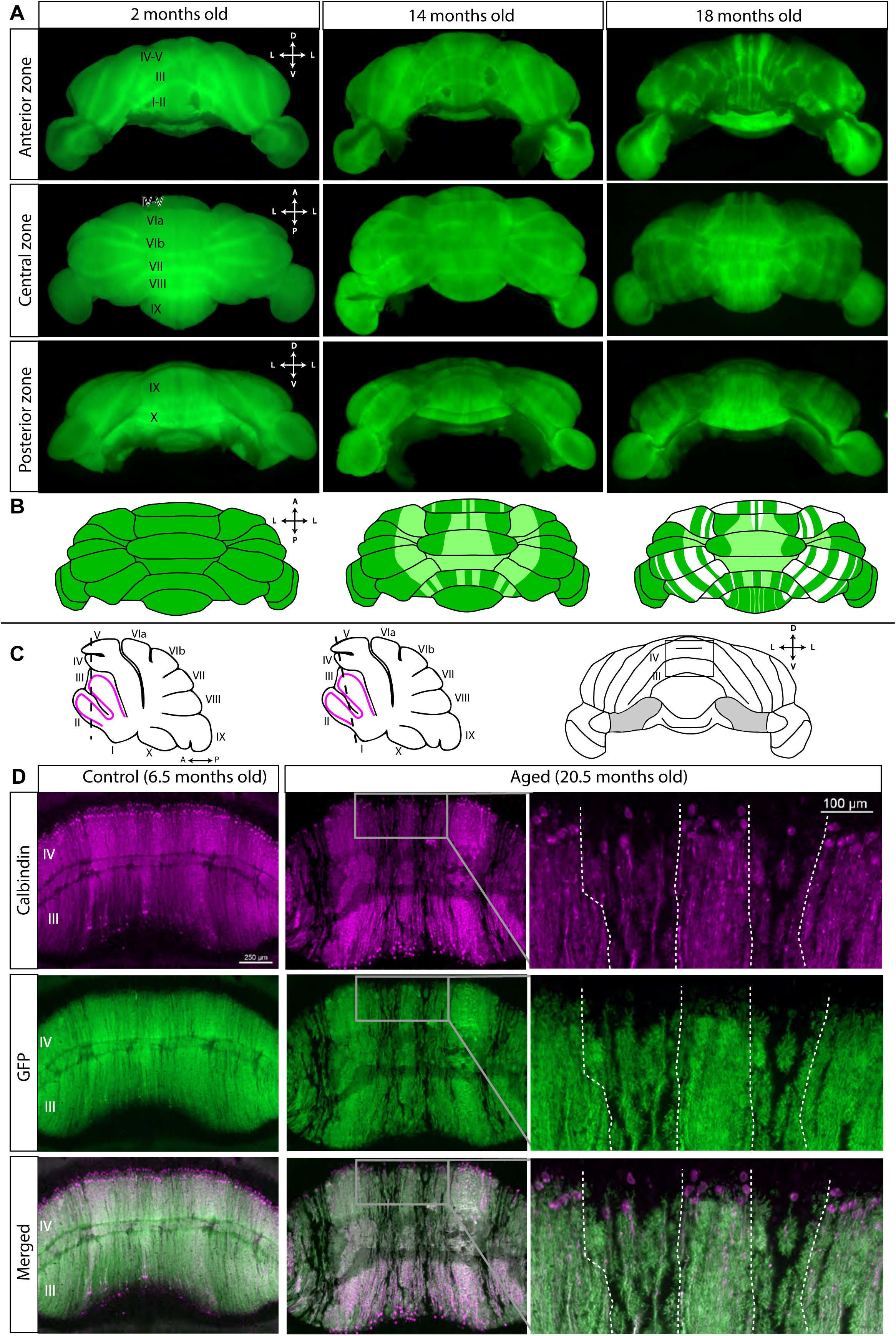
Cerebella of aged Purkinje cell-specific fluorescent reporter mice display the same pattern of Purkinje cell loss as revealed by wholemount calbindin immunohistochemistry. A) Wholemount cerebella of Purkinje cell-specific fluorescent reporter mice visualized with blue light and viewed from different angles. Cerebellar lobules are labeled with Roman numerals. D = dorsal; L = lateral; V = ventral; A = anterior; P = posterior. B) Schematics of the dorsal view of cerebella from young, middle-aged, and older Purkinje cell-specific fluorescent reporter mice. Lighter colors indicate less intense reporter expression. C) Schematics of sagittal sections of the cerebellum and a wholemount cerebellum indicating the location of tissue sections. Dashed lines indicate the position and angle of tissue sections. D) Coronal cut cerebellar tissue sections of Purkinje cell-specific fluorescent reporter mice immunostained for calbindin and GFP. Dashed lines indicate boundaries between surviving Purkinje cells and degenerating Purkinje cells. Scale bar = 250 μm; inset scale bar = 100 μm.

A combination of dynamic calbindin expression and staining artifacts can affect the visualization of Purkinje cells even in the absence of Purkinje cell loss. We observed striped calbindin expression in both young and aged C57Bl/J6 control mice (Supplementary Fig. 4A), possibly due to calcium dynamics. We confirmed that these stripes were not due to widespread Purkinje cell loss by staining adjacent tissue sections with Neutral Red, which revealed that the Purkinje cell bodies were still present (Supplementary Fig. 4A). In addition, in young and aged cerebellar tissue from C57Bl/J6 mice, the Purkinje cell dendrites span the molecular layer (Supplementary Fig. 4A and B), whereas in degenerating tissue, shrunken dendrites, thickened axons, and torpedoes can be observed (Fig. 2A and B). Previous studies have shown that calbindin mRNA and protein are reduced in the cerebella of aged mice, rats, and humans^110,111^, and reduced calbindin immunoreactivity was observed in surviving Purkinje cells in aged rats and in patients with spinocerebellar degeneration^112,113^. Therefore, we argue that calbindin expression alone is not a reliable, sufficient indicator of Purkinje cell loss and should be supplemented with other histological and labeling techniques. Staining artifacts can also give the false appearance of Purkinje cell absence, but background staining or Neutral Red can reveal the Purkinje cells (Supplementary Fig. 4B). For this reason, we used multiple methods to visualize Purkinje cell degeneration and loss. Degenerative Purkinje cell pathology, multiple antibodies, Neutral Red staining, and a Purkinje cell-specific genetically driven fluorescent reporter confirmed that the striped cellular pattern we observed indicates the robust presence of regional Purkinje cell loss in aged mice and that it arises due to neurodegeneration in a subpopulation of Purkinje cells.

### The pattern of age-related Purkinje cell loss overlaps with but is distinct from the overall pattern of zebrin II expression

In mutant mice with Purkinje cell loss (*leaner* and *tottering*^114^*, nervous*^115^*, BALB/c npc^nih^*^116^, *C57BLKS/J spm*^116^, *Cacna1a* null^117^, and acid sphingomyelinase knockout (ASMKO) mice^118^), as well as mice with global brain ischemia^119^, neurodegeneration occurs according to the expression of zebrin II^120^. Zebrin II (an antigen on the aldolase C protein^121^) is the most well-studied cerebellar patterning marker. In vertebrates, zebrin II expression reveals a striking pattern of parasagittal stripes across the cerebellar cortex^91^. Degeneration respects two main populations in the map as revealed by zebrin II expression; for example, in the *nervous* mutant mouse, Purkinje cell loss occurs selectively in zebrin II-positive Purkinje cells^115^, whereas in models of Niemann-Pick type C disease, zebrin II-negative Purkinje cells die first^116^. Aged mice displayed three distinct stripes of surviving Purkinje cells in the anterior vermis (Fig. 1G-I), a pathology that resembles the pattern of zebrin II expression in young mice. Therefore, we asked whether the pattern of age-related Purkinje cell loss and survival indeed reflects the pattern of zebrin II expression. To test this, we cut coronal cerebellar tissue sections from aged mice (n=5) and immunostained them for calbindin (to label surviving Purkinje cells) and zebrin II (to reveal Purkinje cell stripe patterning), using cerebellar tissue from young mice (n=4) as controls.

In the anterior zone, age-related Purkinje cell loss respects the zebrin II boundaries. For example, in lobules III and IV, Purkinje cells began to degenerate selectively in zebrin II-negative stripes P1- and P2-, while the zebrin II-positive stripes remained intact (Fig. 4B). In this region, calbindin and Purkinje cell-specific reporter expression both reflect the same pattern of Purkinje cell loss with respect to zebrin II (Supplementary Fig. 5). In other regions, Purkinje cell loss forms clear stripes despite uniform zebrin II expression; for example, in lobule VI and anterior lobule VII, which are almost entirely zebrin II-positive, Purkinje cells degenerate in stripes (Fig. 4B). This is in contrast to lobule VIII in the same mouse, which is unaffected by Purkinje cell degeneration despite the striped zebrin II expression in this lobule (Fig. 4B). Interestingly, we observed differences in the relationship between age-related Purkinje cell loss and zebrin II expression within a single lobule. In dorsal lobule IX, Purkinje cell loss could occur in zebrin II-negative stripes, whereas in ventral lobule IX, Purkinje cell loss occurs in the medial zebrin II-positive stripe P1+ (Fig. 4B). The paraflocculi display bands of Purkinje cell loss (Fig. 4D), but given the heterogeneity of this region, with its developmentally defined Purkinje cell clusters and its stripes of intensely and weakly zebrin II-expressing Purkinje cells, and its complicated morphology^122,123^, more detailed analysis is required to fully understand the relationship between this pattern and zebrin II expression. Taken together, these data show that although the pattern of age-related Purkinje cell loss can correspond with zebrin II expression –for example, in the anterior zone –the underlying pattern that dictates Purkinje cell loss during normal aging is more complicated than a single stripe marker would indicate. Instead, differential vulnerability to age-related neurodegeneration may result from complex interactions between Purkinje cell lineage, gene expression patterns, and specific functional properties. These distinct patterns in aged mice may uncover previously unidentified subsets within uniform zebrin II areas or provide clues into afferent fiber-to-Purkinje cell functional interactions that could influence long-term circuit function and health.

**Figure 4:**
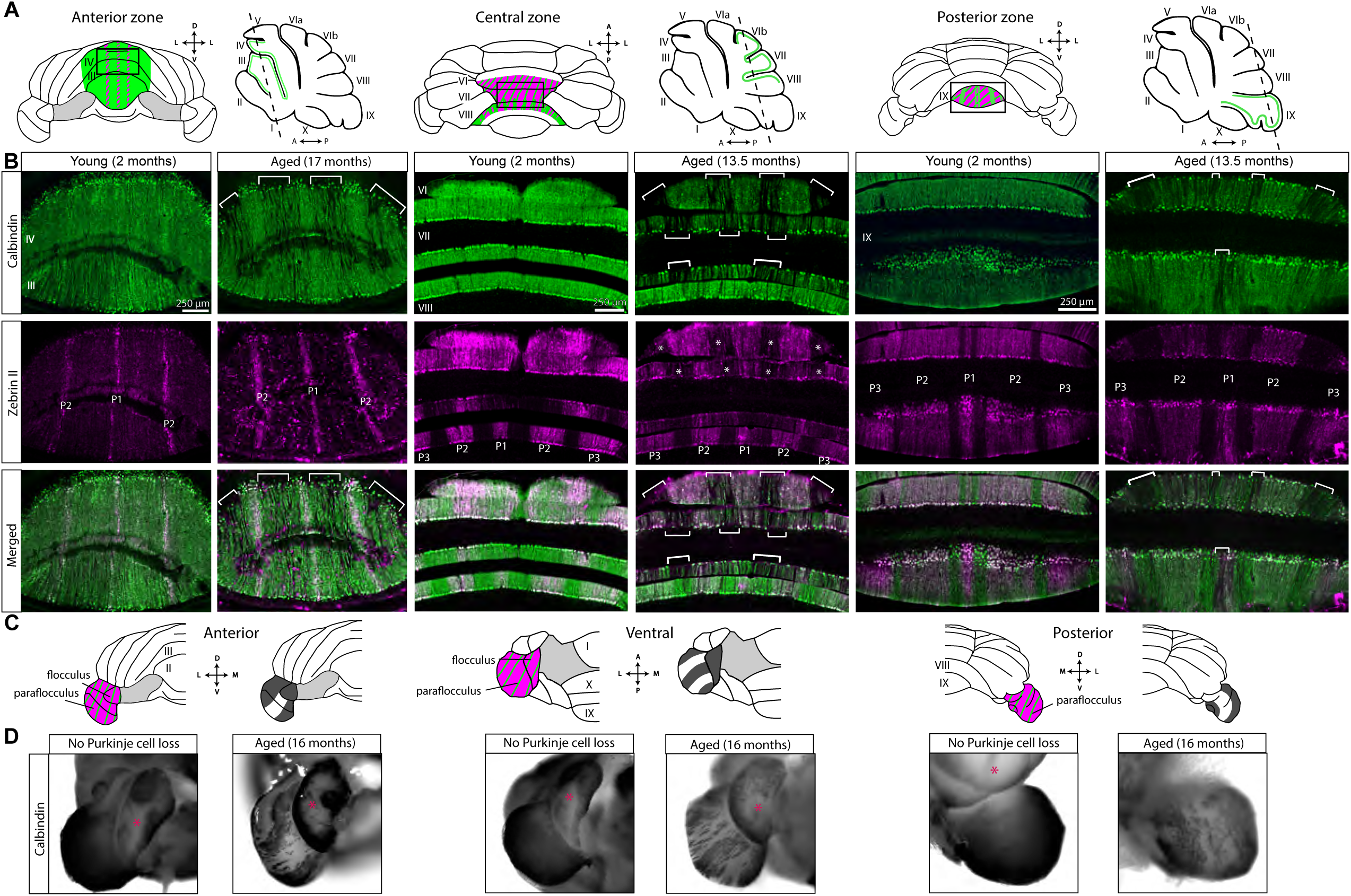
The pattern of age-related Purkinje cell loss has some similarities to zebrin II expression, but the unique overall map represents a greater cerebellar complexity. A) Schematics of wholemount cerebella and sagittal sections of the cerebellum indicating the location of tissue sections. Green indicates calbindin expression and alternating green and magenta indicates where calbindin and zebrin II are co-expressed. Dashed lines indicate the position and angle of tissue sections. Cerebellar lobules are labeled with Roman numerals. B) Coronal cut tissue sections co-stained for calbindin (green) and zebrin II (magenta). Zebrin II-positive stripes are indicated by P1, P2, and P3. Brackets indicate bands of degenerating Purkinje cells, and asterisks indicate bands of degenerating Purkinje cells in uniformly zebrin II-positive regions. Scale bar = 250 μm. C) Schematics of half of a wholemount cerebellum indicating calbindin and zebrin II expression (alternating green and magenta) and bands of surviving Purkinje cells as indicated by calbindin expression (dark gray). D) Paraflocculi of wholemount cerebella immunostained for calbindin and viewed from different angles. Cerebellar lobules are labeled with Roman numerals. D = dorsal; L = lateral; M = medial; V = ventral; A = anterior; P = posterior.

Even during extreme Purkinje cell loss, with few Purkinje cells surviving throughout the cerebellum, a pattern of parasagittal stripes remains visible. This was evident in serial sections taken from a 25-month-old mouse and immunostained for calbindin, revealing the subsets of Purkinje cells that were most resistant to degeneration (Fig. 5). The tissue sections displayed the same pattern of three parasagittal stripes in vermal lobules II through VI, though the width of the stripes was reduced, sometimes to one or two Purkinje cells per stripe. Most of Crus 1, the flocculi, and the paraflocculi had strong calbindin staining, indicating the presence of surviving Purkinje cells. Ventral lobule IX and lobule X were strikingly well preserved in comparison to the rest of the cerebellum. Even within regions with surviving Purkinje cells, the cells were undergoing degeneration. High magnification images of tissue from the 25-month-old mouse revealed extreme morphological abnormalities in Purkinje cells. Beaded recurrent axon collaterals formed plexuses where Purkinje cell somata likely used to be (Fig. 6A-E), and Purkinje cell dendrites were thickened and fractured (Fig. 6A and C). We also observed a putative recurrent axon collateral that extended to the top of the molecular layer in a large gap between surviving Purkinje cells (Fig. 6E). These morphological abnormalities may represent the last efforts of surviving Purkinje cells to reside in an aged cerebellum, where the majority of Purkinje cells have degenerated.

**Figure 5:**
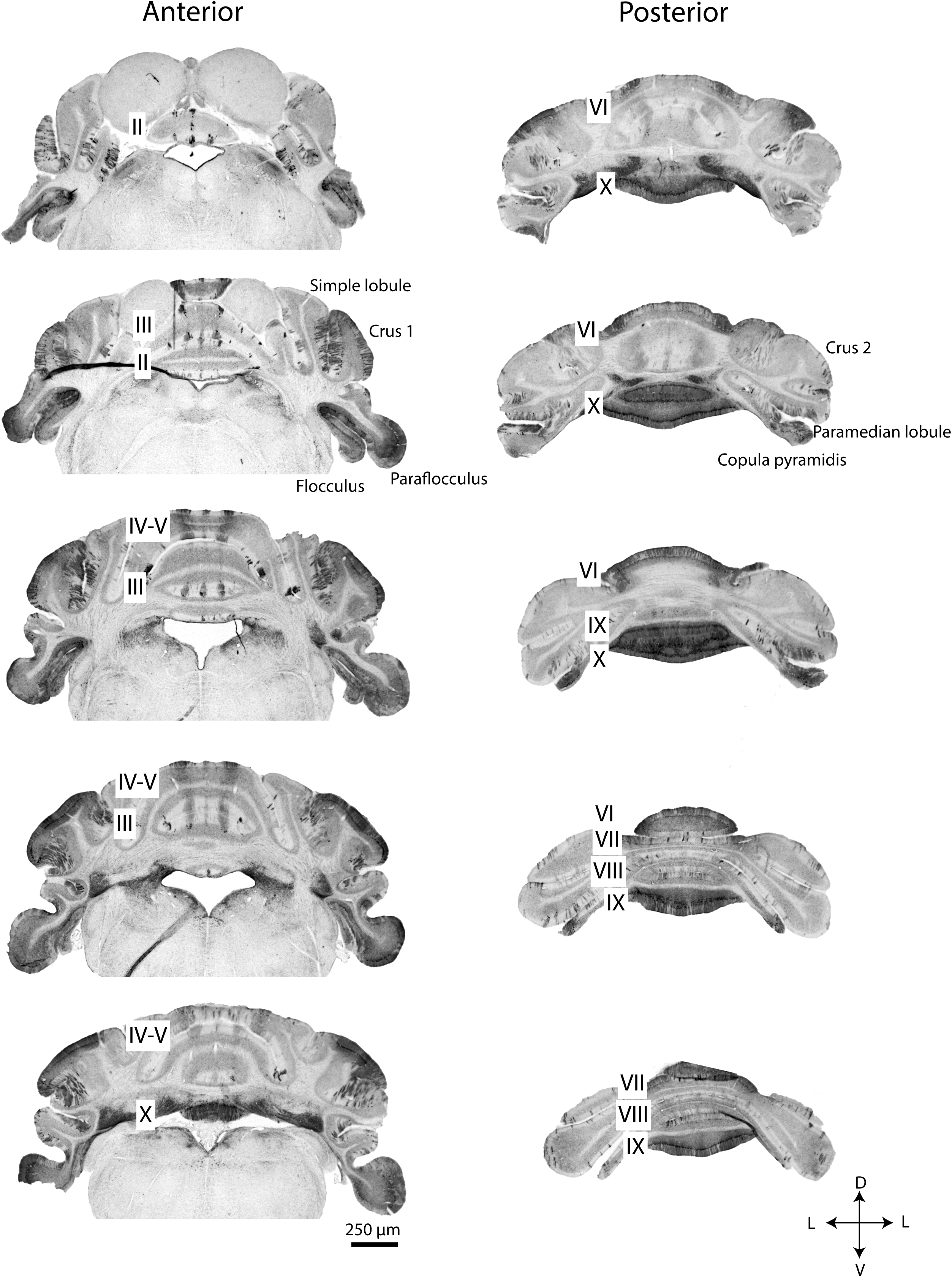
Regions with lasting resistance to Purkinje cell loss during normal aging are revealed in serial sections from 25-month-old mouse. Coronal cut cerebellar tissue sections immunostained for calbindin and arranged in order from anterior to posterior. Cerebellar lobules are labeled with Roman numerals. D = dorsal; L = lateral; V = ventral; A = anterior; P = posterior. Scale bar = 250 μm.

**Figure 6:**
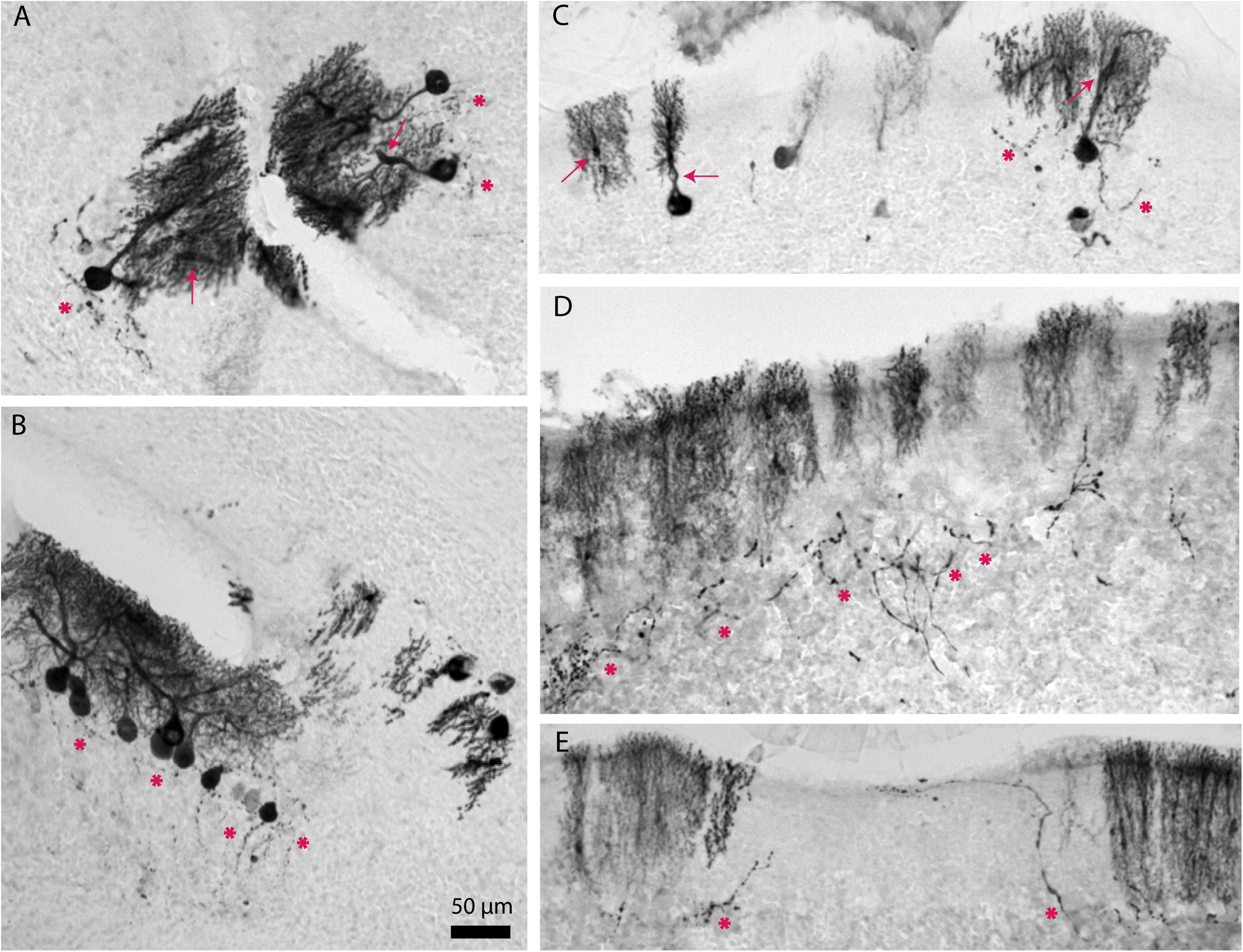
Regions with apparent resistance to cell loss in 25-month-old mouse have Purkinje cells with extreme morphological abnormalities. High-magnification images of cerebellar tissue sections immunostained for calbindin. Asterisks indicate recurrent axon collaterals, and arrows indicate thickened dendrites. Scale bar = 50 μm.

### Light sheet imaging reveals a pattern of age-related Purkinje cell loss in the cerebellum

Light sheet imaging, when combined with tissue clearing, allows the visualization of labeled cells in multiple dimensions throughout a brain structure. Given the precise regional specificity of age-related Purkinje cell loss, we performed light sheet imaging of the cerebellum of an aged Purkinje cell-specific fluorescent reporter mouse (Video 1) to fully visualize the pattern. Light sheet imaging revealed that lobule X is resistant to Purkinje cell loss during normal aging, similar to our observations of wholemount cerebella immunostained for calbindin. Lobule X, the flocculi, and the paraflocculi comprise the nodular zone, a largely zebrin II-positive region (Fig. 4C). The paraflocculi showed stripes of Purkinje cell loss during aging, as seen on calbindin-stained wholemounts (Fig. 4D). By combining the advantages of a Purkinje cell-specific fluorescent reporter with the ability to reveal patterns within the core of the cerebellum (which are typically hard to appreciate due to the folding of the cortex), where antibodies do not always penetrate, light sheet imaging provides a complete picture of the pattern of age-related Purkinje cell loss.

Taken together, our results from the wholemount cerebella, coronal tissue sections, and light sheet imaging of Purkinje cell-specific reporter expression indicate that despite some similarities, the pattern of age-related Purkinje cell loss is similar, but not identical to the expression pattern of zebrin II, a reliable marker that defines the endogenous map of Purkinje cell stripes and zones.

### Motor function is impaired in aged mice compared to young mice

Our results show that Purkinje cell loss during normal aging is extensive and occurs throughout the cerebellum. Despite this, the general motor behavior of our cohort of aged mice was ostensibly normal when the mice were observed in their home cages. To investigate the effect of age-related Purkinje cell loss on motor behavior more closely, we performed a series of behavioral tests, including the accelerating rotarod, the horizontal ladder, and analysis in a tremor monitor, on young and aged mice before sacrificing them and histologically examining the cerebellum for Purkinje cell loss. We found that the aged mice (n=12) had a shorter latency to fall from the accelerating rotarod compared to the young mice (n=8; Fig. 7B). Although the latency to fall was always lower in the aged mice compared to the young mice, the aged mice improved from day to day (Fig. 7B), demonstrating their continued capacity and ability to learn new motor skills.

**Figure 7:**
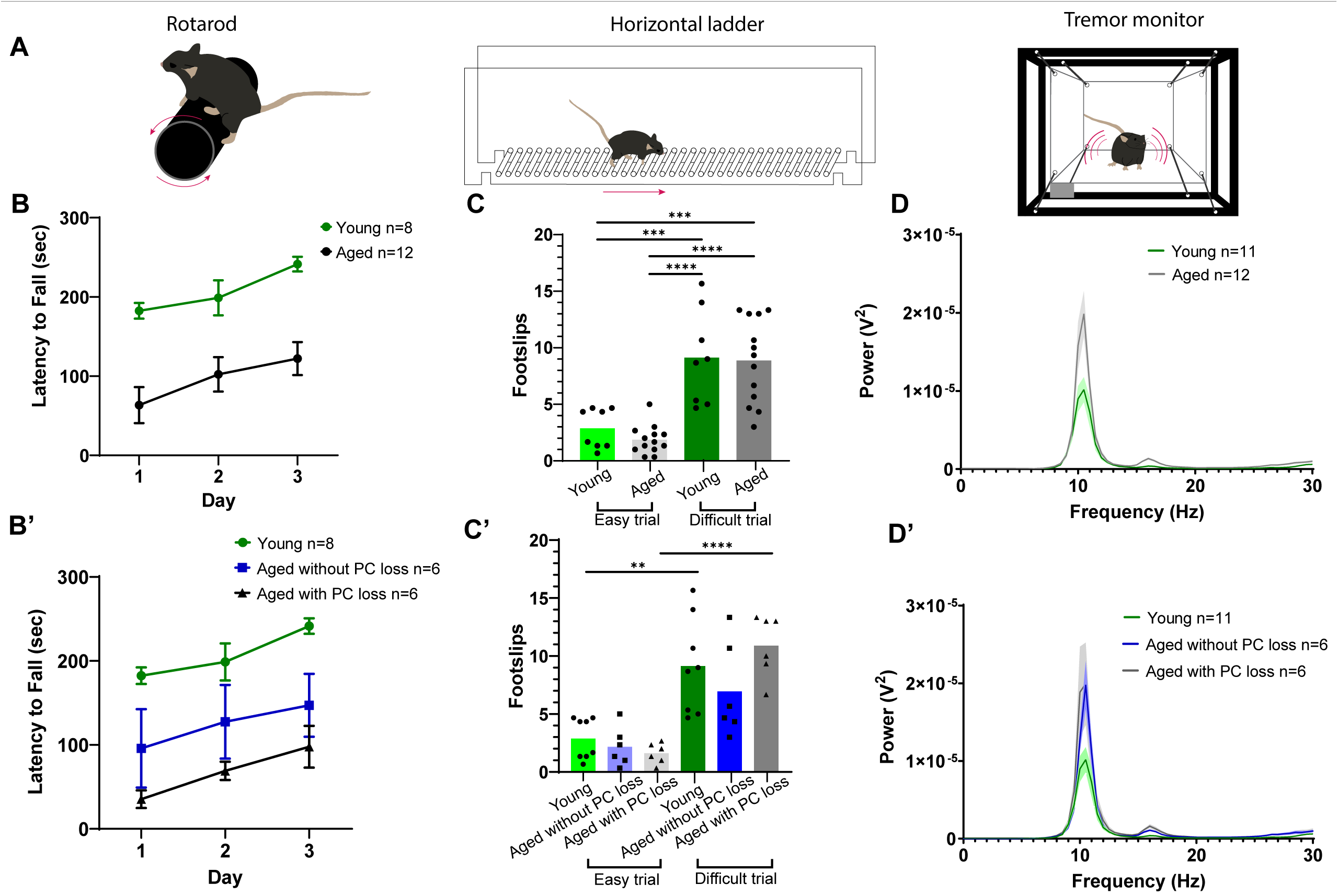
Aged mice display impaired performance on the accelerating rotarod and increased tremor but no deficits on the horizontal ladder. A) Schematics of motor function tests. B) Latency to fall from accelerating rotarod. Error bars indicate standard error of the mean. B’) Latency to fall with aged mice sorted based on the presence or absence of Purkinje cell loss. C) Number of footslips when crossing the horizontal ladder. ** indicates p ≤ 0.01, *** indicates p ≤ 0.001, and **** indicates p ≤ 0.0001. C’) Number of footslips with aged mice sorted based on the presence or absence of Purkinje cell loss. D) Power spectrum of tremor detected by a tremor monitor. Error bars indicate standard error of the mean. D’) Power spectrum of tremor with aged mice sorted based on the presence or absence of Purkinje cell loss.

To test skilled, voluntary movement and limb control, we used a horizontal ladder task. Mice were subjected to an “easy” trial, where every ladder rung was in place, and a “difficult” trial, where every other ladder rung was removed. The number of footslips was recorded per trial. As expected, both the young (n=8) and aged (n=12) groups had more footslips during the difficult trial compared to during the easy trial (Fig. 7C). However, we did not detect a significant difference in the number of footslips between the young and aged groups during either trial (Fig. 7C).

Because humans and mice have increased tremor with age^124,125^ and since the cerebellum is implicated in tremor pathophysiology^17,19,21^, we used a custom-built tremor monitor to quantify tremor power and frequency^126^. We found that aged mice (n=12) had significantly higher tremor power compared to young mice (n=11; Fig. 7D). Peak power in both cohorts occurred at a frequency of ∼ 10 Hz (Fig. 7D), consistent with instances of pathological tremor being found within the frequency range of physiological tremor in mice^125,126^.

We next wondered what structural and/or functional variables might contribute to the motor deficits observed in aged mice, using peak tremor power as an example. We tested whether peak power in aged mice (n=12) was influenced by body weight, relative age, or sex. Female aged mice tended to weigh less and have lower power tremor but still overlapped with male aged mice in terms of weight and peak tremor power (Supplementary Fig. 6A). We did not find a statistical correlation between peak tremor power, weight, or relative age (defined as the age of an individual mouse in comparison to other mice in the aged group; Supplementary Fig. 6B). This suggests that although aged mice have significantly increased tremor power compared to young mice (Fig. 7D), neither weight nor differences in relative age among aged mice contribute significantly to tremor within the aged group. In other words, tremor does not necessarily worsen with increased age beyond a certain point within a given age range. This may be related to our finding that middle aged mice can have Purkinje cell loss while some older mice do not (Table 1). Thus, aging is not a simple linear process in which increasing age is always negatively (or gradually) correlated with the loss of specific neural circuit functions and a decline in specific behaviors.

Given that not all aged mice have striped Purkinje cell loss, even within the same litter (Fig. 1D-I; Table 1), we wondered whether aged mice without Purkinje cell loss performed better on motor tasks compared to aged mice with Purkinje cell loss. To address this, after behavioral testing, we collected cerebellar tissue sections from the aged mice and immunostained them with calbindin antibody. We subdivided the behavioral data of the aged mice based on whether they had Purkinje cell loss or not. We found that there was no significant difference in rotarod performance, number of footslips on the horizontal ladder, or tremor between aged mice with Purkinje cell loss and aged mice without Purkinje cell loss, though both aged groups had shorter latency to fall on the rotarod and an increased tremor power compared to young mice (Fig. 7B’, C’, and D’). Together, these results suggest that while aged mice exhibit abnormalities in tremor and motor coordination, Purkinje cell loss alone, and specifically mild Purkinje cell loss, may not cause these behavioral impairments. Alternatively, Purkinje cell dysfunction, to varying degrees, may set a platform for the development of tremor and motor incoordination in aging mice, which could then co-initiate different abnormal behaviors with a given amount Purkinje cell loss.

### Postmortem tissue from neurologically normal humans reveals age-related Purkinje cell degeneration with the co-presence of healthy and pathological cells

Reports that cerebellar volume is differentially impacted across lobules in human aging^47,55,61,76–79^ prompted us to examine human cerebellar tissue at the cellular level to determine whether Purkinje cells exhibit differential vulnerability and resistance to age-related degeneration. We studied postmortem cerebellar tissue from three neurologically normal patients: a 21-year-old, a 57-year-old, and a 74-year-old. Using the calbindin antibody that effectively and reliably labels Purkinje cells in mice, on sagittal cut tissue sections through the vermis, we observed well-preserved Purkinje cells with expansive dendritic arbors in the tissue from the 21-year-old (Fig. 8). In contrast, the immunostained tissue from the 74-year-old revealed extensive Purkinje cell loss that was observed as noticeable gaps in the Purkinje cell layer that were devoid of somata. In addition, we observed Purkinje cell dendrites with poor integrity and less span in their typically expansive architecture, a change that is indicative of retraction and degeneration of Purkinje cell dendrites (Fig. 8), similar to our observations of Purkinje cells in aged mice (Fig. 2A and B). Interestingly, the tissue from the 57-year-old represented an “intermediate” stage with many robust healthy Purkinje cells that were flanked by areas with Purkinje cell dendrite deterioration and gaps in the Purkinje cell layer (Fig. 8). These data imply the presence of progressive degeneration and loss of Purkinje cells during human aging, as well as differential vulnerability and resistance to such loss among individual cells.

**Figure 8:**
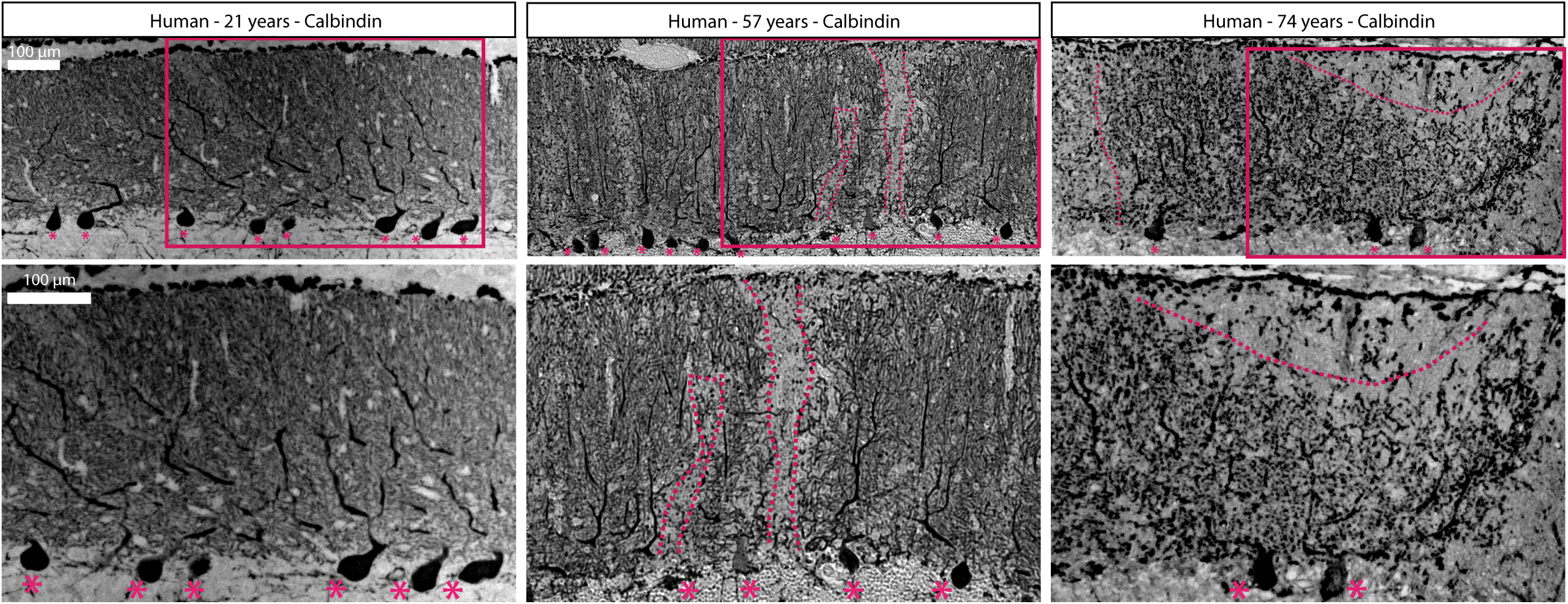
Aging humans have Purkinje cell degeneration that can be visualized with calbindin. Asterisks indicate the remaining Purkinje cell bodies. Dashed lines indicate the boundaries of Purkinje cell dendritic arbors. Scale bar = 100 μm.

## DISCUSSION

We demonstrate that Purkinje cell loss that occurs during normal aging is not uniform. Instead, Purkinje cell loss occurs in a striking pattern of parasagittal stripes in aged mice. While this striped pattern bears some resemblance to the pattern of zebrin II expression, especially in the anterior zone lobules, the overall pattern is different from zebrin II expression, which typically defines the known stripe patterns of Purkinje cell loss in disease models. Furthermore, we show that aged mice have increased tremor power and deficits in rotarod performance compared to young mice but that performance on the horizontal ladder is preserved. Overall, we have found that age-related Purkinje cell loss occurs in a distinct striped pattern that may provide insight into the selective vulnerability and resistance of cells to neurodegeneration in the normal aging cerebellum.

### Striped Purkinje cell loss during normal aging is more complex than zebrin II expression

Although evidence of striped Purkinje cell loss has not been reported in aged mice prior to our study, striped Purkinje cell loss is widely appreciated in mouse models of disease. In disease models, Purkinje cell loss typically occurs preferentially in either zebrin II-positive (e.g. *nervous* mutant mice^115^) or zebrin II-negative (e.g. *BALB/c np^nih^* and *C57BLKS/J spm* mice^116,118^) Purkinje cells^120^. Interestingly, we found that although age-related Purkinje cell loss occurred preferentially in zebrin II-negative stripes in the anterior zone, it followed a unique region-to-region pattern in the rest of the cerebellar cortex. Similar findings, in which Purkinje cell loss occurs in zebrin II-negative stripes in the anterior zone but not the posterior zone, have been observed in a mouse model of autosomal-recessive spastic ataxia of Charlevoix-Saguenay (ARSACS)^127^, yet the pattern of age-related Purkinje cell loss is distinct. This suggests that other factors in addition to zebrin II identity may influence the selective vulnerability to neurodegeneration. In addition, neurodegeneration is a continuous process. The loss of certain Purkinje cells may trigger the degeneration of nearby cells regardless of regional vulnerability, creating a domino effect that may not precisely reflect zonal markers.

Importantly, zebrin II stripes are not the only example of cerebellar molecular patterning, and therefore interpretation of patterned Purkinje cell loss should not be limited to the pattern of zebrin II expression alone. There are many molecular markers whose expression patterns form stripes in Purkinje cell subpopulations that correspond with the zebrin II stripes (e.g. GABA-B receptor^83^ and PLCβ3^82^), are complementary to zebrin II stripes (e.g. PLCβ4^82^), or have a more complex relationship (e.g. HNK1^128^, HSP25^129^, and NFH^84^). In multiple rodent models with Purkinje cell loss, Purkinje cells in the nodular zone that express HSP25 are more resistant to degeneration than HSP25-negative Purkinje cells^116,118,129^. In addition to molecular markers, cerebellar zones are also defined by their developmental lineage, climbing fiber and mossy fiber inputs, interneuron organization, and Purkinje cell outputs to the cerebellar nuclei^98^, any of which might contribute to differential Purkinje cell vulnerability. Therefore, the pattern of Purkinje cell loss during normal aging may reflect a previously unidentified zone marker or a combination of pattern modalities.

Furthermore, we report differences in the susceptibility to Purkinje cell loss not only between Purkinje cell subpopulations but between individual mice. This is evident given that some mice, even mice as old as 18-24 months, do not display Purkinje cell loss (Table 1). These aged mice without Purkinje cell loss do not exhibit Purkinje cell pathology indicative of neurodegeneration, although more subtle morphological changes may occur that escape our detection. Especially intriguing is our finding that there are littermate pairs in which one littermate has striped Purkinje cell loss, and the other littermate does not. This effect may not be dictated solely by genotype or sex (Table 1). Variable phenotypes among littermates is not unusual – for example, graying hair, hair loss, differences in energy levels, posture, metabolism, weight, and body fat content –and this variability can increase with age^130^. It is possible that the presence or absence of age-related striped Purkinje cell loss is another such variable phenotype. This would present an interesting scenario given the striking difference between a cerebellum with no apparent Purkinje cell loss and a cerebellum with broad stripes of degenerated Purkinje cells; because the pattern adheres to a strict organizational code, this may result in a more binary readout than other metrics that vary with age. One factor that could contribute to this variability is animal housing density. Group housed mice are more variable in their body compositions compared to singly house mice^131^, and group and single housing can each result in different behavioral stressors (aggression among males and social isolation, respectively)^132,133^. These stressors can have behavioral and physiological effects, including neural and endocrine changes^132,133^, and may indirectly contribute to differential Purkinje cell vulnerability and resilience between individuals during aging.

### A combination of methods confirms patterned Purkinje cell degeneration and/or cell loss

Here, we report the presence of reproducible, striped Purkinje cell loss in aged mice, whereas previous studies of aged rodents have concluded that Purkinje cell loss is largely uniform throughout the cerebellum^62–66^. This discrepancy may be due to the limitations of using tissue sections alone to study patterned neurodegeneration. In previous studies, quantification of age-related Purkinje cell loss has been performed by counting cells in sagittal sections of the cerebellum taken every 320 μm^63–66^. To use the pattern of Purkinje cell loss observed in our study as an example, the observations would vary greatly depending on where a sagittal section was taken. For example, a sagittal section taken at the midline could reveal little or no Purkinje cell loss, whereas a section taken more laterally could reveal large swathes of missing Purkinje cells. The full stripes could only be properly and fully appreciated with coronal sections, which risks leaving out information about anterior-posterior dendrite defects that can be better appreciated with sagittal sections. Therefore, relying on only coronal or sagittal sections is likely insufficient to appreciate complex regional differences in Purkinje cell vulnerability during aging. Our study has the advantage of using wholemount visualization and light sheet imaging in addition to coronal and sagittal tissue sections, enabling the multi-dimensional visualization of cerebellar patterning.

Methods-based discrepancies in regional cerebellar degeneration have also been observed in elderly patients. One imaging study found reduced volume in the hemispheres and vermal lobules VI-X while the anterior vermis was unaffected^47^, but a later study by the same authors found uniform vulnerability across the vermis and attributes the differences to methodological differences^48^, underscoring the importance of factoring technique into results about regional degeneration in the highly compartmentalized cerebellum. However, the majority of studies in elderly patients have found that when cerebellar atrophy is observed, atrophy is most severe in the anterior cerebellum^55–57,61,77,79^, with significant volume reduction or Purkinje cell loss often observed in the vermis^55,58,76,78^. Similarly, we found that aged mouse cerebella with striped Purkinje cell loss had the most profound loss in the anterior cerebellum, including the vermis (Fig. 1A). This similarity in morphological phenotype, combined with our finding of Purkinje cell loss in cerebellar tissue from middle-aged and older, neurologically normal human patients (Fig. 8), suggests translatability between our findings of Purkinje cell loss in aged mice and humans. However, a much larger study must be undertaken with a greater number of human tissue specimens, accounting for a wider span of ages, possible gender differences, race, and other person to person variabilities such as socioeconomic status, diet, exercise, and overall health. To determine the extent to which our findings in mice relate to human aging, it will also be important to study regional Purkinje cell loss at the cellular level in elderly humans. As the field currently stands, studies of cerebellar atrophy in humans lack the resolution to examine Purkinje cell degeneration at the cellular level, and detailed analysis of human Purkinje cell subpopulations is confounded by the size and complexity of the cerebellum. Studies of human postmortem cerebellum tissue have long neglected the complex organization of the cerebellum, but a recent study used a marker of Purkinje cell stripes in mice to differentiate Purkinje cell subtypes in human tissue, thereby identifying differential axonal pathology in essential tremor^134^. A similar method could be useful for studying the link between patterned Purkinje cell loss in mice and humans.

### Regional vulnerability creates distinct patterns of cerebellar degeneration in diseases versus during normal aging

In humans, cerebellar degeneration occurs in distinct patterns reflected by the affected lobules in a given subtype of neurodegenerative disease^135^ and whether the patient is affected by a neurodegenerative disease or normal aging^77,136^. The pattern of cerebellar degeneration may impact the manifestation of symptoms because of the functional compartmentalization of the cerebellum at the level of lobules and the stripes within these lobules. Purkinje cell synaptic plasticity and firing rates differ depending on the stripes they inhabit^137^, and modules differ in terms of behavioral function, although these functions likely overlap across modules to some degree^138^. This suggests that cerebellar compartments and Purkinje cell subpopulations may represent a key for unlocking the neural correlates of cerebellar dysfunction in disease and aging.

The mechanisms underlying patterned cerebellar degeneration remain unknown, though evidence points to the differential vulnerability of Purkinje cell subpopulations^119,129,139–141^. One potential mechanism for differential Purkinje cell vulnerability is stripe-specific excitotoxicity. Our study found that in aged mice, where Purkinje cell loss corresponds with zebrin II expression, Purkinje cell loss occurs preferentially in zebrin II-negative stripes. Zebrin II-negative Purkinje cells have a higher average firing frequency than zebrin II-positive Purkinje cells^142,143^, which may make zebrin II-negative Purkinje cells more susceptible to excitotoxicity. Accordingly, excitatory amino acid transporter 4 (EAAT4) expression is restricted to zebrin II-positive Purkinje cells, and deafferentation of Purkinje cells prevents ischemia-induced Purkinje cell loss in zebrin II-negative stripes^119^. This selective loss of Purkinje cell subpopulations during aging has functional implications. Recent work has shown that Purkinje cells in aged mice have a reduced firing rate compared to Purkinje cells in young mice^144^. Furthermore, the distribution of Purkinje cell firing frequencies is similar in young and aged mice, but higher-firing Purkinje cells are absent in aged mice^144^, suggesting that higher-firing Purkinje cells die while lower-firing cells survive. This corresponds with the results of our study, in which higher-firing zebrin II-negative Purkinje cells tend to degenerate before lower-firing zebrin II-positive Purkinje cells during aging. Stripe-specific excitotoxicity, in combination with potentially neuroprotective proteins such as HSP25^145^, may influence differential Purkinje cell vulnerability in aged mice. Understanding the mechanistic origins of differential vulnerability and resistance in distinct cell populations could shed light on therapeutic methods to block or even prevent neurodegeneration in disease and aging.

### Mild age-related Purkinje cell loss alone may not cause overt motor impairments

The aged mice in our study displayed overtly normal motor behavior during observation, while behavioral tests revealed that aged mice had higher power tremor than young mice and impaired performance on the rotarod. These findings are in accordance with previous behavioral studies of normal aged mice^74,125^. Interestingly, aged mice performed as well as young mice on the horizontal ladder test, even when the difficulty was increased by removing every other ladder rung (Fig. 7C). This may be because aged mice can adapt abnormal kinematics for voluntary movements to compensate for gradual functional deficits during the aging process, whereas involuntary, whole-body tasks such as the accelerating rotarod prove more difficult to overcome.

Upon examination of the cerebella, the aged mice used in these behavioral tests had either no Purkinje cell loss or mild Purkinje cell loss with no evident stripes (severe cases would typically be accompanied by a clear striped pattern). To determine the influence of Purkinje cell loss on age-related motor deficits, we separated the behavioral data based on the presence or absence of Purkinje cell loss. We did not find a significant difference in motor behavior between the two aged subgroups, which suggests that mild Purkinje cell loss alone is not sufficient to cause the observed motor deficits. The aging process involves the dysfunction and degeneration of different classes of neurons within different brain regions (with both overlapping and distinct temporal onsets), which likely all co-contribute to the decline of motor behavior over time. Additionally, it remains to be determined whether the Purkinje cells in aging mice are functionally normal before their degeneration. It is possible that deterioration begins earlier and that they are contributing to network dysfunction well before their eventual degeneration and that all aged mice are vulnerable to motor deficits when Purkinje cells display abnormal physiological properties. In-depth, precise behavioral phenotyping over time will be necessary to investigate a potential link between progressive changes in Purkinje cell morphology and motor function during aging.

We focused this study on mice of the C57BL/6J background. A next step would be to investigate age-related Purkinje cell loss in other inbred and outbred strains. Different strains, including C57BL/6J, display differences in motor behavior tests^146,147^, as well as differences in cerebellar morphology^148^. Therefore, whether strain-specific patterns of Purkinje cell loss and motor behavior exist in aged mice is an important question for future research.

## CONCLUSION

We show that Purkinje cell loss occurs in a striped pattern during normal aging. We revealed this patterned neurodegeneration using a combination of wholemount immunohistochemistry, tissue staining on sections, and transgenic mice encoded with a Purkinje cell-specific reporter. Our work establishes a fresh perspective for how patterns of degeneration in models of aging and disease could inform about symptomology and regional vulnerability. Despite the apparent chaos of widespread Purkinje cell degeneration, the strict organization of the cerebellum established early in development lends an order to the chaos. Future studies will benefit from identifying the causes of differential vulnerability across Purkinje cell subpopulations and between individual mice, as well as investigating Purkinje cell loss and motor function in aged mice of multiple strains. By understanding the mechanisms of patterned, age-related Purkinje cell loss, we would better appreciate the functional implications of neurodegeneration during normal aging. Eventually, the characteristics that confer resistance to neurodegeneration in specific Purkinje cell subpopulations may prove useful for designing effective treatments that maximize features of healthy brain aging.

## MATERIALS AND METHODS

### Mice

Mouse husbandry and experiments were performed under an approved Institutional Animal Care and Use Committee (IACUC) protocol at Baylor College of Medicine (BCM). All mice used in this study were maintained in our colony at BCM. Mice between 2 and 5 months of age were categorized as young mice, 11-15 months old were considered middle-aged, and older than 16 months were considered old (consistent with information at Jackson Laboratory). Both males and females were used. A mixed population of mice was used, some of which were C57BL/6J mice ordered from Jackson Laboratory (#000664) and some of which were multiple generations descended from the following Jackson Laboratory strains: #006207, #007900, #012882, #014647, #024948, #12569, #12897, #21162, #024109, #021188, and #24109. No conditional knockout or knock-in crosses were used except for *Pcp2^Cre^;Ai32* and *Pcp2^Cre^;Ai40D* mice, which were used to fluorescently tag Purkinje cells. *Pcp2^Cre^;Ai32* and *Pcp2^Cre^;Ai40D* mice were used interchangeably and will be referred to as Purkinje cell-specific fluorescent reporter mice throughout the study.

### Perfusion and sectioning

Mice were anesthetized by intraperitoneal injection of Avertin (2, 2, 2-Tribromoethanol, Sigma-Aldrich catalog #T4). Cardiac perfusion was performed with 0.1 M phosphate-buffered saline (PBS; pH 7.4), followed by 4% paraformaldehyde (4% PFA) diluted in PBS. Brains were dissected and post-fixed at 4°C for at least 24 hr in 4% PFA. Brains were then cryoprotected in sucrose solutions (10% sucrose in PBS, then 20% sucrose, then 30% sucrose) and embedded in Tissue-Tek O.C.T. compound (Sakura Finetek USA; catalog #4583). Tissue sections were cut on a cryostat with a thickness of 40 μm and placed in PBS.

### Free-floating tissue section immunohistochemistry

Immunohistochemistry on free-floating frozen-cut tissue sections was performed as described previously^149^. Rabbit anti-calbindin (1:10,000; Swant) or mouse anti-calbindin (1:2,000; Sigma) was used to label all Purkinje cells. Mouse anti-zebrin II (1:250; gift from Dr. Richard Hawkes, University of Calgary, Alberta, Canada) was used to reveal Purkinje cell stripes. Immunoreactive complexes were visualized with either 3,3’-diaminobenzidine tetrahydrochloride (DAB, 0.5 mg/mL; Sigma-Aldrich), nickel-DAB (DAB Substrate Kit; Vector Labs), or anti-mouse or anti-rabbit secondary antibodies conjugated to fluorophores (1:1,500; Invitrogen). For DAB reactions, horseradish peroxidase (HRP)-conjugated goat anti-rabbit and goat anti-mouse secondary antibodies (diluted 1:200 in PBS; DAKO) were used to bind the primary antibodies. Antibody binding was revealed by incubating the tissue in the DAB solution, which was made by either dissolving a 100 mg DAB tablet in 40 mL PBS and 10 μL 30% H_2_O_2_ or using the DAB Substrate Kit. The DAB reaction was stopped with PBS when the optimal color intensity was reached.

After staining, tissue sections were placed on electrostatically coated glass slides. Tissue sections were coverslipped using either Cytoseal (for DAB) or FluoroGel with Tris buffer (for immunofluorescence; Electron Microscopy Sciences).

### Neutral Red

After tissue sections were placed on electrostatically coated glass slides, they were left to dry overnight. Then, slides were dipped briefly in distilled water before being immersed in 1% Neutral Red for 30 min. Next, slides were subjected to an ethanol series and xylene before being coverslipped with Cytoseal.

### Wholemount immunohistochemistry

Wholemount immunohistochemistry was performed as previously described^100,101^. Cerebella were post-fixed in Dent’s fixative for 6 hr at room temperature (RT) and bleached in Dent’s bleach overnight at 4°C. The next day, cerebella were dehydrated in two 30-min rounds of methanol (MeOH) at RT. The cerebella were then subjected to five freeze/thaw cycles before being placed at −80°C overnight. The next day, the cerebella were rehydrated with washes in 50% MeOH/50% PBS, 15% MeOH/85% PBS, and 100% PBS for 1 hr each at RT. The tissue was enzymatically digested with 10 ug/mL Proteinase K in PBS for 3 min. Then, the cerebella were washed in PBS three times for 10 min each at RT. Tissue was blocked in PBSMT overnight at 4°C. The next day, the tissue was incubated in primary antibodies diluted in PBSMT with 5% DMSO for 48 hr at 4°C. After incubation, the cerebella were washed in PBSMT twice for 2 hr each at 4°C. Then, the tissue was incubated in secondary antibodies diluted in PBSMT with 5% DMSO at 4°C overnight. The next day, the cerebella were washed in PBSMT twice for 2 hr each at 4°C, followed by a single wash in PBT for 1 hr. Finally, the cerebella were incubated in DAB solution for 10 min. The DAB reaction was stopped by placing the cerebella in 0.04% sodium azide.

### Microscopy

Images of stained tissue sections were captured with either a Zeiss AxioCam MRm (fluorescence) camera mounted on a Zeiss Axio Imager.M2 microscope or a Leica DMC2900 (brightfield) camera mounted on a Leica DM4000 B LED microscope. Wholemount cerebella immunostained for calbindin were placed in 1% agar and immersed in PBS, then imaged with a Zeiss AxioCam MRc 5 camera mounted on a Zeiss Axio Zoom.V16 microscope. Brains from Purkinje cell-specific fluorescent reporter mice were imaged with a Zeiss AxioCam MRm camera mounted on a Zeiss Axio Zoom.V16 microscope immediately after perfusion. After imaging, the raw data was imported into Adobe Photoshop, which was used to correct brightness and contrast levels. Schematics were created in Adobe Illustrator.

### Clearing and light sheet imaging

Brains were cleared with the EZ Clear method as described previously^150^. Whole brain images were acquired with a Zeiss Light Sheet Z1 at a refractive index of 1.52 with a 5x objective. Image tiles were stitched together with Stitchy and visualized with Arivis.

### Accelerating rotarod

The rotarod (ENV-571M, Med Associates, Inc., Vermont, USA) was set to accelerate from 4 to 40 rpm in 5 min (setting 9) and was stopped at 300 sec if the mice successfully stayed on for this duration. Mice rested for at least 10 min between trials. Rotarod performance was measured in three trials per day for three consecutive days.

### Tremor monitor

Tremor was measured using a custom-built model similar to that described previously^126^. Mice were placed in a translucent plastic box with an open top. The box was held steady in the air by eight elastic cords attached to corners of the box and to a scaffold. An accelerometer at the bottom of the box detected movements of the box. Signals from the accelerometer were recorded and analyzed in Spike2 software. Power spectrums of tremor across naturally occurring tremor frequencies (0-30 Hz) were made using a fast Fourier transform (FFT) with a Hanning window. An offset was applied to center the tremor waveform on 0, and the recordings were down sampled to produce frequency bins aligned to whole numbers. For each mouse, the first 120 sec of recording in the tremor monitor was defined as the acclimation period, and the following 180 sec of recording was used for analysis.

### Horizontal ladder

The horizontal ladder test was performed on a custom-built ladder consisting of rods placed horizontally between two plexiglass walls. Mice were placed at the entrance of the ladder and allowed to walk across. If a mouse turned around before completing the test, the mouse was placed back at the entrance of the ladder to restart the trial. Video recordings of each trial were analyzed to count the number of foot slips per 50-cm section of ladder. A foot slip was counted when a foot passed below the level of the rungs. The easy horizontal ladder test involved rods spaced 1 cm apart, and the difficult test involved the removal of every other rod, resulting in rods spaced 2 cm apart. Each mouse completed three trials of the easy test in one day, followed by three trials of the difficult test the next day.

### Human tissue

Use of human postmortem brain tissue was granted exemption by the Baylor College of Medicine Institutional Review Board. All procedures involving a human participant were performed in accordance with the National Research Committee and the 1964 Declaration of Helsinki and its later amendments or comparable ethical standards. The three brains were removed as part of routine hospital autopsy with no significant neurological history or neuropathological findings. Age was extracted from the autopsy report.

### Quantitative analyses and statistics

Fisher’s exact test was performed to determine whether the presence of striped Purkinje cell loss in aged mice is sex dependent. Mice were grouped by sex and by whether striped Purkinje cell loss was observed.

Pixel intensity of calbindin and GFP was calculated with the plot profile function in ImageJ (version 1.54g). Plot profile was performed on identically sized rectangular regions of interest that were placed over lobules II and III. One anterior coronal tissue section immunostained for calbindin or calbindin plus GFP was analyzed per mouse. Because pixel intensity is a measure of brightness, immunofluorescence has a high pixel intensity, whereas the DAB chromogen has a low pixel intensity. For this reason, pixel intensity was used to plot mediolateral differences in immunofluorescence, and inverse pixel intensity was used to plot mediolateral differences on tissue sections where DAB was used to visualize the staining.

Molecular layer thickness was calculated by measuring the molecular layer in dorsal lobule VIII in coronal tissue sections immunostained with calbindin (DAB). We chose to measure lobule VIII because of its striking zebrin II pattern and because the cerebellar cortex of this lobule has regions without curvature that are ideal for measuring. One tissue section was used per mouse, and three measurements of the molecular layer were taken per tissue section (midline, left of midline, and right of midline). The three measurements in each tissue section were averaged, and the averages were plotted. The data was analyzed with one-way ANOVA with multiple comparisons.

For the accelerating rotarod and horizontal ladder tests, the three trials per day were averaged, and the averages were plotted. The horizontal ladder data was analyzed with one-way ANOVA with Tukey’s multiple comparisons test.

## ACKNOWLEDGEMENTS

This work was supported by Baylor College of Medicine, Texas Children’s Hospital, the Jan and Dan Duncan Neurological Research Institute (Texas Children’s Hospital Duncan NRI), the National Institute of Neurological Disorders and Stroke (RVS: R01NS119301 and R01NS127435; SGD: F31NS129279), Eunice Kennedy Shriver National Institute of Child Health and Human Development of the National Institutes of Health under Award Number P50HD103555 for use of the Cell and Tissue Pathogenesis Core (the BCM IDDRC), and the BCM Optical Imaging and Vital Microscopy Core, with the expert assistance of Chih-Wei Hsu. Roy V. Sillitoe is supported by the Ting Tsung and Wei Fong Chao Foundation. The content is solely the responsibility of the authors and does not necessarily represent the official views of the National Center for Research Resources or the National Institutes of Health.

## ETHICS

Animal experimentation: Mice were housed in an AAALAS-certified animal facility. All procedures to maintain and use these mice were approved by the Institutional Animal Care and Use Committee for Baylor College of Medicine (Animal protocol number AN-5996).

## CONTRIBUTIONS

Sarah G. Donofrio, Cheryl Brandenburg, Amanda M. Brown, and Roy V. Sillitoe contributed to the study conception and design. Material preparation, data collection, and analysis were performed by SGD, CB, AMB, and RVS. HCL provided human postmortem tissue, and TL processed and stained the human postmortem tissue. The first draft of the manuscript was written by SGD, SGD and RVS wrote and assembled the full manuscript, and all authors commented on each version of the manuscript. All authors read and approved the final manuscript.

## SUPPLEMENTARY xxFIGURES AND xxVIDEOS

**Supplementary Figure 1: The presence of Purkinje cell loss varies across aged mice.**

A) Proportion of aged mice with striped Purkinje cell loss, non-striped Purkinje cell loss, and no Purkinje cell loss divided by sex. B) Mediolateral inverse pixel intensity of Purkinje cell labeling in aged mice. Inverse pixel intensity was calculated within identically sized rectangular regions of interest placed over lobules II and III in coronal cut tissue sections immuostained for calbindin. Data were grouped according to whether mice had non-striped Purkinje cell loss (blue) or striped Purkinje cell loss (black). B’) Mediolateral inverse pixel intensity of Purkinje cell labeling separated by individual aged mouse.

**Supplementary Figure 2: Different calbindin antibodies reveal matching expression patterns, including age-related striped Purkinje cell loss.** Coronal tissue sections taken from the anterior and posterior cerebellum of young and aged mice and co-immunostained with two different calbindin antibodies. Scale bar = 500 μm.

**Supplementary Figure 3: Calbindin and GFP reveal overlapping patterns of expression in aged mice with striped Purkinje cell loss.** A) Mediolateral pixel intensity of Purkinje cell labeling in aged mice. Pixel intensity was calculated within identically sized rectangular regions of interest placed over lobules II and III in coronal cut tissue sections taken from aged Purkinje cell-specific fluorescent reporter mice and co-immunostained for calbindin and GFP. Calbindin pixel intensity is shown in magenta, and GFP pixel intensity is shown in green. A’) Mediolateral pixel intensity of Purkinje cell labeling separated by individual aged mice.

**Supplementary Figure 4: Patterned calbindin expression and staining artifacts can obscure Purkinje cell loss due to neurodegeneration.**

A) Coronal cut tissue sections from C57BL/6J mice either immunostained for calbindin alone or immunostained for calbindin and stained with Neutral Red. Brackets indicate regions of zonal calbindin expression. Calbindin zones, when present, arise due to domains of high versus low expression rather than extended areas that completely lack calbindin-expressing Purkinje cells. Dashed lines indicate boundaries formed by intact Purkinje cell dendrites and degenerating dendrites. Arrowheads indicate calbindin-negative Purkinje cell bodies that are stained with Neutral Red. Cerebellar lobules are labeled with Roman numerals. Scale bar = 250 μm; inset scale bar = 100 μm. B) Coronal cut tissue sections from aged C57BL/6J mice either immunostained for calbindin alone or immunostained for calbindin and stained with Neutral Red. Arrowheads indicate Purkinje cell bodies that were not stained with calbindin antibody. Scale bar = 100 μm; inset scale bar = 50 μm.

**Supplementary Figure 5: Calbindin and Purkinje cell-specific fluorescent reporter expression reflect the same stripes of Purkinje cell loss with respect to zebrin II.** Coronal cut tissue sections taken from lobules II-III of young and aged Purkinje cell-specific fluorescent reporter mice and co-immunostained for GFP, zebrin II, and calbindin. Scale bar = 250 μm.

**Supplementary Figure 6: There is no correlation between weight, peak tremor power, or relative age.** A) Peak tremor power plotted against body weight, with point color indicating age and point size indicating sex. B) Correlation matrix of peak tremor power, body weight, and age calculated using Spearman’s correlation.

**Video 1: Light sheet imaging reveals the pattern of age-related Purkinje cell loss throughout cerebellum.** Light sheet imaging through the fluorescent reporter cerebellum reveals the pattern of cell loss with excellent anatomical continuity from its surface features to its internal architecture.

