## Supplementary figures and images for "Cerebellar Purkinje cell stripe patterns reveal a differential vulnerability and resistance to cell loss during normal aging in mice"

### Supplementary Figure 1

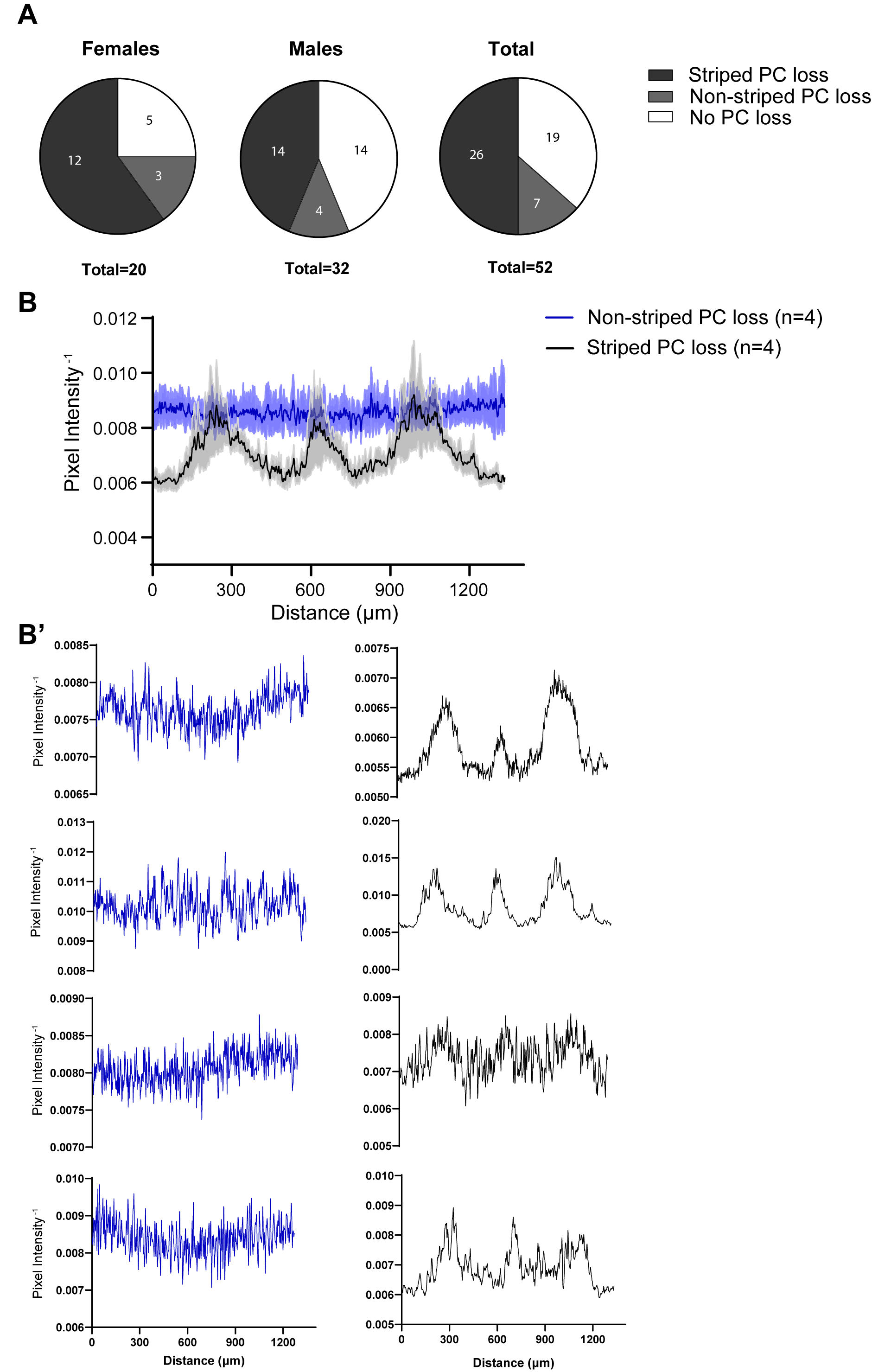

### Supplementary Figure 2

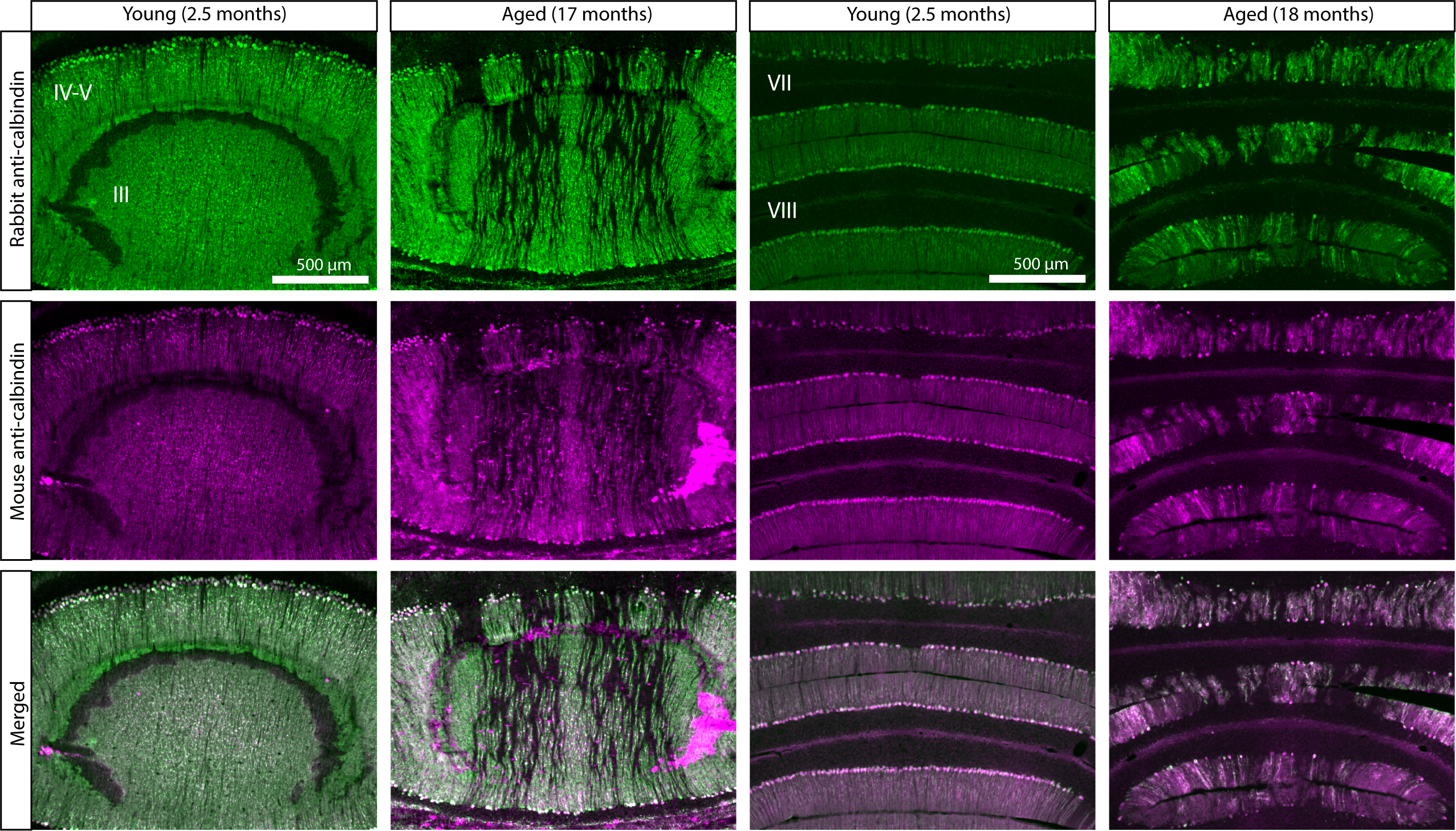

### Supplementary Figure 3

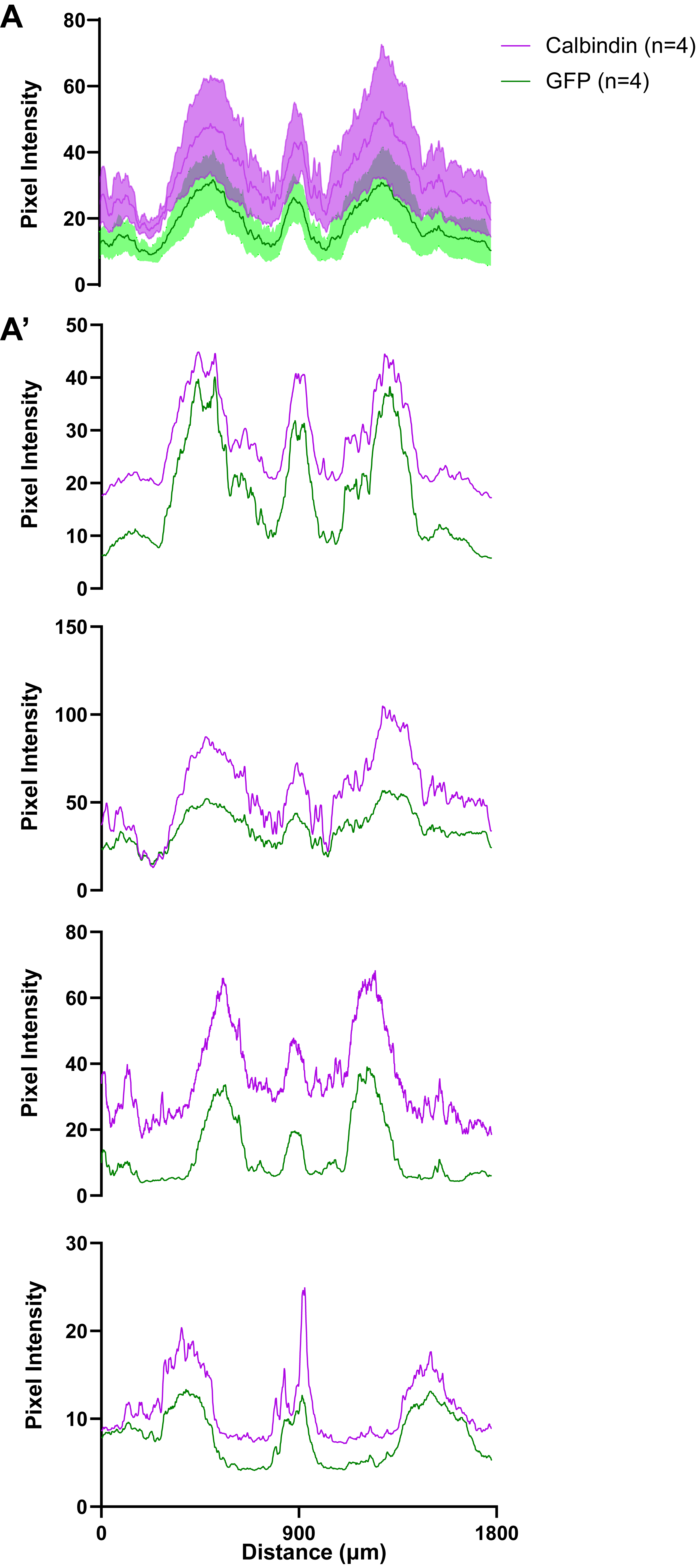

### Supplementary Figure 4

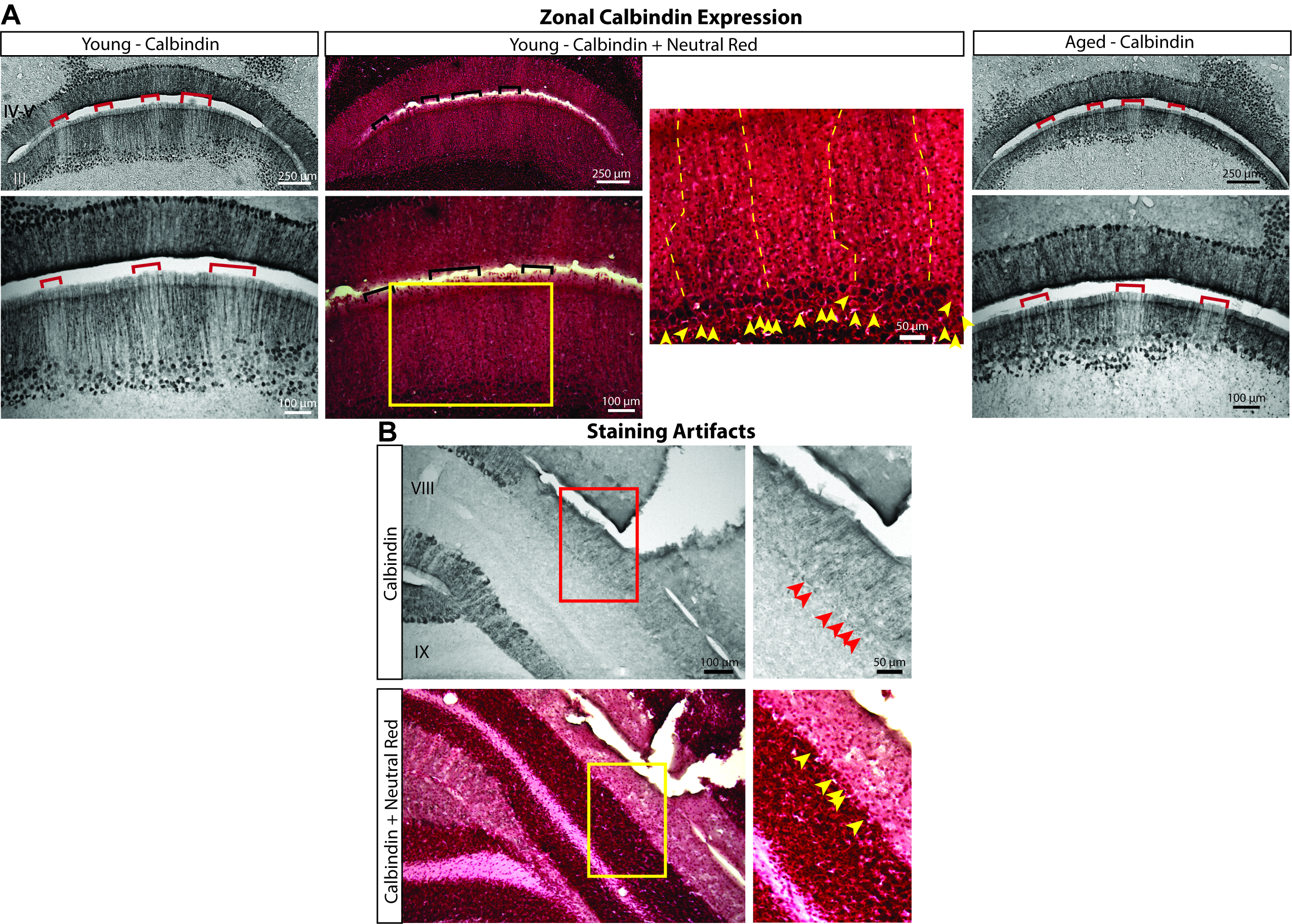

### Supplementary Figure 5

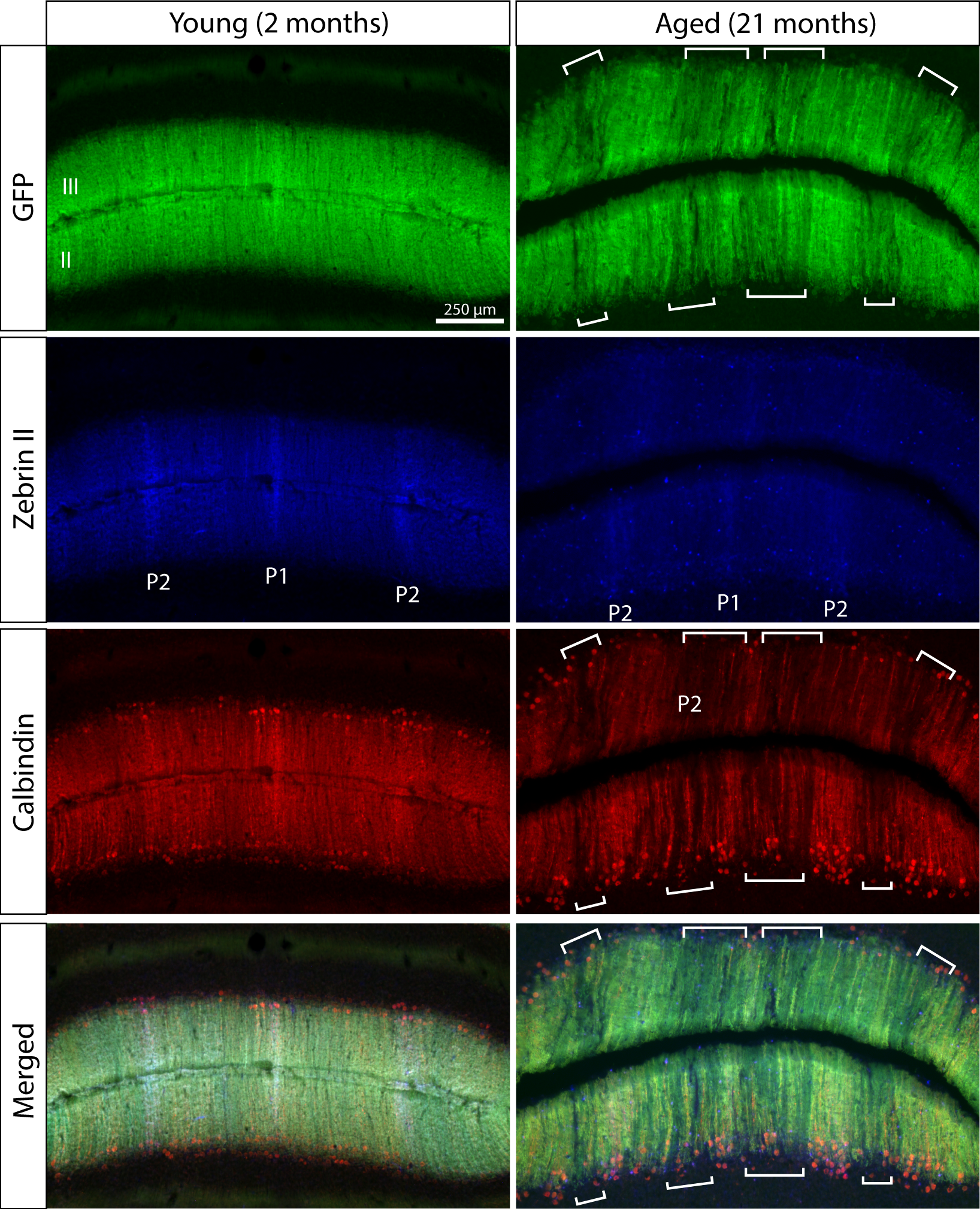

### Supplementary Figure 6

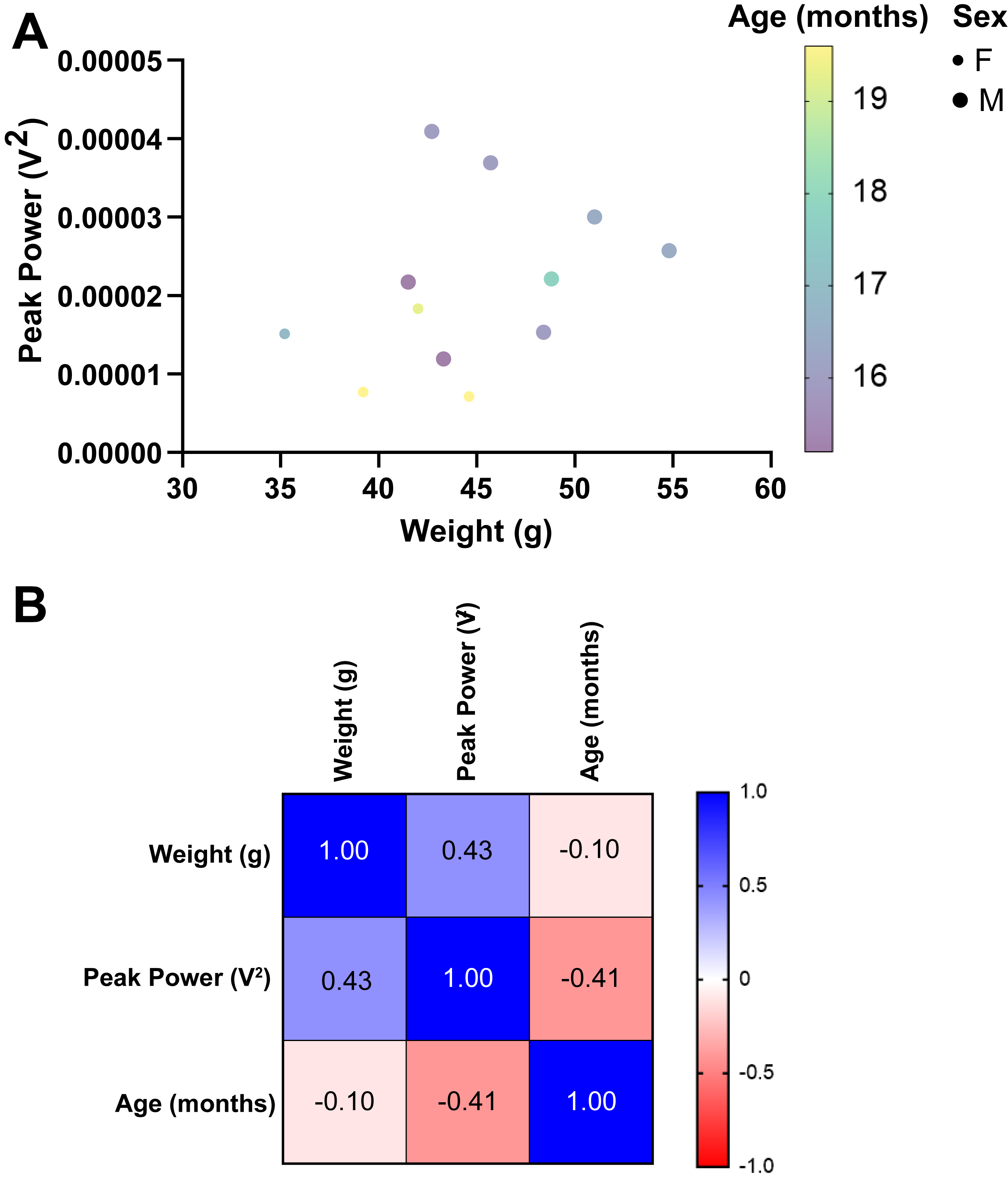
